# Robustness of Felsenstein’s versus Transfer Bootstrap Supports with respect to Taxon Sampling

**DOI:** 10.1101/2023.02.27.530178

**Authors:** Paul Zaharias, Frédéric Lemoine, Olivier Gascuel

## Abstract

The bootstrap method is based on resampling alignments and re-estimating trees. Felsenstein’s bootstrap proportions (FBP) is the most common approach to assess the reliability and robustness of sequence-based phylogenies. However, when increasing taxon-sampling (i.e., the number of sequences) to hundreds or thousands of taxa, FBP tends to return low supports for deep branches. The Transfer Bootstrap Expectation (TBE) has been recently suggested as an alternative to FBP. TBE is measured using a continuous transfer index in [0,1] for each bootstrap tree, instead of the {0,1} index used in FBP to measure the presence/absence of the branch of interest. TBE has been shown to yield higher and more informative supports, without inducing falsely supported branches. Nonetheless, it has been argued that TBE must be used with care due to sampling issues, especially in datasets with high number of closely related taxa. In this study, we conduct multiple experiments by varying taxon sampling and comparing FBP and TBE support values on different phylogenetic depth, using empirical datasets. Our results show that the main critic of TBE stands in extreme cases with shallow branches and highly unbalanced sampling among clades, but that TBE is still robust in most cases, while FBP is inescapably negatively impacted by high taxon sampling. We suggest guidelines and good practices in TBE (and FBP) computing and interpretation.

---

Branch supports are essential for interpreting trees because they allow quantifying the degree of uncertainty in our phylogenetic hypotheses. For Maximum Likelihood (ML) tree estimation (and other approaches, e.g. distance-based), the most popular branch support is undeniably the bootstrap method proposed by Felsenstein (1985), one of the most cited article of all time (Van Noorden et al. 2014). The procedure relies on resampling with replacement the sites of a reference alignment until obtaining a pseudo- (or bootstrap) alignment of same length. Then, pseudo- (or bootstrap) trees are estimated using the same inference method. Finally, the support for every branch on the reference tree is measured as the bootstrap proportion (BP) of pseudo-trees containing that branch. The interpretation of bootstrap support values has led to great controversies in the 1990s (reviewed in Sanderson 1995; Soltis and Soltis 2003; Simon 2020). Originally, Felsenstein (1985) suggested to interpret it as a measure of repeatability, meaning the “probability that a specified group will be found in an analysis of an independent sample of characters” (Hillis and Bull 1993). However, this interpretation has been contrasted with another view that bootstrap could be interpreted as a confidence region of some kind in a null hypothesis framework (Hillis and Bull 1993), and was further discussed in the literature (Felsenstein and Kishino 1993; Sanderson 1995; Efron et al. 1996; Susko 2009) but didn’t find success among practitioners. Through simulations, Hillis and Bull (1993) suggested a threshold value of 70%, although this value was proposed under very specific conditions, namely “equal rates of change”, “symmetric phylogenies”, and “internodal change of <20% of the characters”. Despite those important limitations, Soltis and Soltis (2003) note that “many systematists have adopted Hillis and Bull’s “70%” value as an indication of support”, an observation still true today, even though the “95%” and the more arbitrary “80%” cut-offs can often be seen in the literature.

Aside from conceptual matters, two main criticisms against FBP are recurring in the literature, especially for “large” datasets. By large datasets, we mean in this study matrices where the number of taxa is large, while the number of sites remains moderate, usually corresponding to a single gene or phylogenetic marker. The first is technical: re-estimation (usually by ML) of pseudo-trees is computationally demanding on large datasets and can be impractical on very large datasets. A Rapid Bootstrap Support (RBS; Stamatakis et al. 2008) was proposed and consists in using some of the trees found during the ML topological research as bootstrap trees. The RBS is fast and reliable, but it remains computationally intensive and tends to be more liberal than standard FBP (Anisimova et al. 2011). Even faster is the Ultrafast Bootstrap approximation approach (UFBoot; Minh et al. 2013; Hoang et al. 2018), but recent works suggest it is considerably more liberal than standard FBP (and RBS), questioning to the comparability of UFBoot to standard bootstrap (Gascuel and Lemoine 2022). The second main criticism is FBP’s sensitivity to rogue taxa (Wilkinson 1996), that is, taxa whose position varies from (pseudo-)trees to (pseudo-)trees. Indeed, if a single taxon is unstable (e.g., due to homoplasy or missing data) in the overall tree or in a particular region of it, then the FBP support values are expected to be considerably lowered in that region. Other criticisms discuss the problem of “large” dataset in the sense of “large number of sites” (e.g., Sharma and Kumar 2021) but this will not be covered here as it is not the subject of this study.

The commonly used alternatives to Felsenstein Bootstrap Support (FBP) can be divided into two main classes. The first one is the Posterior Probability (PP) of Bayesian phylogenetics (Rannala and Yang 1996), calculated as the proportion of all sampled trees in the MCMC chain (post burn-in) in which the branch of interest is found. Due to their parametric nature and implementation, PP are often considered liberal, as opposed to the more conservative FBP. Abundant literature exists on comparing BP and PP (e.g., Douady et al. 2003), but expanding on that matter goes beyond the frame of this article. Local supports are another class of alternative supports, responding to a need for computational speed in a context of ever-growing datasets. These supports are obtained by locally rearranging the tree topology around the branch of interest, using Nearest Neighbor Interchange (NNI). Some of the most popular ones include the approximate likelihood-ratio test (aLRT; Anisimova and Gascuel 2006) and the non-parametric Shimodaira–Hasegawa-like version (aLRT SH-like; Guindon et al. 2010). However, these approaches only provide a local view of the support. It is also important to note that none of these two classes explicitly addresses the problem or rogue taxa: Bayesian PP are expected to behave similarly to FBP with a tree that contains rogues; the local supports are little affected by the presence of a few rogues, but also unable to detect them and measure their impact on the overall tree due to their local nature.

Recently, an alternative to FBP has been proposed: the Transfer Bootstrap Expectation (TBE; Lemoine et al. 2018). TBE is also based on resampling and re-estimating pseudo-trees, and thus is also computationally heavy, although easily parallelizable and applicable with RBS and UFBoot. The difference lies in the comparison of the pseudo-trees to the reference tree. Rather than the binary presence/absence of a reference branch in the pseudo-trees, TBE uses a “transfer” distance that is measured using the number of taxa that must be transferred (or removed) to make two branches identical (Lemoine et al. 2018). Because of its continuous nature, TBE scores are always higher than FBP, except for cherries (i.e., a clade comprising only two taxa) where both supports are identical. TBE is also less affected by rogues (Lemoine et al. 2018) and it has a natural interpretation: on a given reference branch, we can easily calculate the average number of taxa that have to be transferred to recover that branch in bootstrap trees. Finally, yet importantly, results with real and simulated data showed that TBE supports do not induce falsely supported branches. While their initial results using simulations and empirical datasets indicate that 70% is a reasonable threshold for supporting branches, Lemoine et al. (2018) suggest that it is “better to interpret TBE values depending on the data and the phylogenetic question being addressed”. TBE also has solid mathematical ground, and is guaranteed to converge in probability to 0 when the size of the tree grows and there is no signal in the data (Dávila Felipe et al. 2019).

TBE was proposed in a context of ever-growing number of sequences, in particular in the epidemiology field. It has become recurrent in the literature to find extremely large datasets (in terms of number of sequences): bacteria/archaea (10,575 tips; Zhu et al. 2019), mushrooms (5,284 tips; Varga et al. 2019), fishes (31,526 tips; Rabosky et al. 2018), diatoms (19,197 tips; Lewitus et al. 2018), angiosperms (36,101 tips; Janssens et al. 2020), and especially viruses like HIV (9,147 tips, Lemoine et al. 2018) and of course SARS-CoV-2 (1.6 million tips; Turakhia et al. 2022). In datasets of this size, it is expected that some of the taxa will behave like rogues, thus lowering FBP values. In fact, it is not rare to find FBP scores below 20% (or even at 0%), especially for deep branches, even when those have strong phylogenetic signal. Yet these deep branches are usually the primary focus of large-scale studies. Lemoine et al. (2018) showed that TBE was in fact able to support deep branches that have strong phylogenetic signal without being affected by a few rogues.

Since its publication five years ago, TBE has been cited about 400 times (Google Scholar, Feb/2023), but has generated little debate. However, a recent review from Simon (2020) expressed a stimulating critic, that “TBE must be used with care due to sampling issues. For example, if many closely related taxa are added to the tree TBE values will increase across the entire tree. This is because the measure is based on counting the number of sequences sampled, not taking into account their variation.” This critic raises a series of important questions related to sampling variation (or taxon sampling) and sampling disequilibrium in large datasets. In this study, we will explore the impact on both FBP and TBE of taxon sampling, rogue taxa and sampling disequilibrium in large datasets, using theoretical examples and empirical datasets. We provide a series of guidelines in the Discussion for good practices and interpretation when estimating FBP and TBE support values on large datasets.

## Theoretical results

In this section, we explore two theoretical cases with sampling variation and presence of rogues. The goal is mainly pedagogical, partly to address the critic published in Simon (2020), but it is unlikely that any of these examples could be found in real data, that we explore further. To facilitate the reader’s comprehension, we remind here the basic formulae for FBP and TBE.

In FBP, the support of a branch (or bipartition) in a tree *T* is the proportion of bootstrap trees *T\** in which the bipartition is present. If a bipartition *b\** from *T\** is found identical as *b* in *T*, then the support of *b* given *T\** is equal to 1, else it is equal to 0. As opposed to the binary nature of FBP, TBE was proposed as a continuous version, using the transfer distance. The distance *δ*(*b,b\**) is equal to the number of taxa that must be transferred (or removed) to make both bipartitions identical. The transfer index *Φ*(*b,T\**) is the minimum of the transfer distances for all bipartitions *b\** present in *T\**: *Φ*(*b,T*^∗^) = *Min*_*b*_∗_∈*T*_^∗^{*δ*(*b, b*^∗^)}. This index is then normalized in [0,1] and averaged over all bootstrap trees. The normalization relies on *p*, the size of the smallest of the two subsets of taxa defined by *b*. It is easily seen that *Φ*(*b,T*^∗^) ≤ *p* − 1. The normalized version of the transfer index is thus equal to 1 − *Φ*(*b,T*^∗^) /(*p* − 1), and the TBE support is defined by: 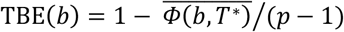, where the bar denotes the average over all bootstrap trees. It follows that TBE(*b*) ≥ FBP(*b*) when using the same set of bootstrap trees. The difference between the two supports depends on *p* and thus the depth of *b* (as in Lemoine et al. 2018, we assume here and in the whole article that the depth of a branch is measured by *p*). If *p* = 2 (i.e., a cherry), then both supports are equal. If *b* is a deep branch and *p* is large, there will often be a large difference between the two supports. This is what we will explore in this study, first through theoretical examples, and then with real data.

### Impact of duplicated sequences on TBE scores

In his personal communication to Simon (2020), Nick Goldman suggests that, for a given phylogenetic signal, TBE supports will be increased if we add many closely related taxa to the tree. A model that closely matches this hypothesis is presented hereafter. Let P|Q be a bipartition of reference tree *T*, where P and Q have sampling size *p* and *q*, respectively, *p* ≤ *q* and *p* + *q* = *n*, where *n* corresponds to the total number of taxa (*p* is again the depth of bipartition/branch P|Q). To keep the same phylogenetic signal while increasing the sampling, we simply add duplicated taxa (i.e., strictly identical sequences). The number of taxa will be multiplied by a factor *k*, where *k* represents the number of duplicated sequences, *k* = 1 corresponds to the original dataset, and *p* and *q* are transformed into *kp* and *kq* when duplicates are added. With any reasonable phylogenetic program, the reference and bootstrap trees will remain essentially the same, with all duplicated sequences grouped together into clusters that form the “tips” of the tree, while the rest of the tree and the internal branches are unchanged. In this model, the phylogenetic signal remains the same and the FBP support is unchanged regardless of the value of *k*.

Let us now describe the behavior of the TBE supports. In the absence of duplicates (*k* = 1), the TBE support of P|Q (let us call it *σ*(1)) is equal to 1 − *τ*/(*p* – 1), where *τ* is the average number of taxa that need to be transferred to retrieve P|Q in the bootstrap trees. Then we have *τ* = (1 − *σ*(1))(*p* − 1). Let us now assume that each sequence is duplicated *k* times. Then, it is easily seen that the support of the duplicated version of P|Q is equal to:

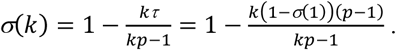

Using the derivative, we note that *σ*(*k*) is an increasing function of *k*. When *k* is large, *σ*(*k*) converges to *σ*_*max*_ = *σ*(1)(1 – 1/*p*) + 1/*p*, and with large *p* we have *σ*(1) ≈ σ_*ma*x_ ≈ *σ*(k) for any value of *k*. In other words, the support of large clades (deep branches) remains unchanged when adding duplicates. However, things are different when *p* is small (i.e., close to 2). Figure 1 shows the TBE supports for different values of *p* and *k*, and for *σ*(1) = 0.3, 0.7 and 0.9 (i.e., very low, moderate and high phylogenetic signal). We see that sampling redundancy (i.e., the addition of duplicates) has a strong impact on cherries (*p* = 2), much less on small clades (*p* = 3, 5, 10), and very little when the clade is large (*p* = 100). The gap between FBP and TBE is maximized for cherries: with *p* = 2, FBP = TBE and FBP is not impacted by duplicates, whereas TBE is clearly impacted and increases with the number of duplicates, especially if the support is low.

**Figure 1:**
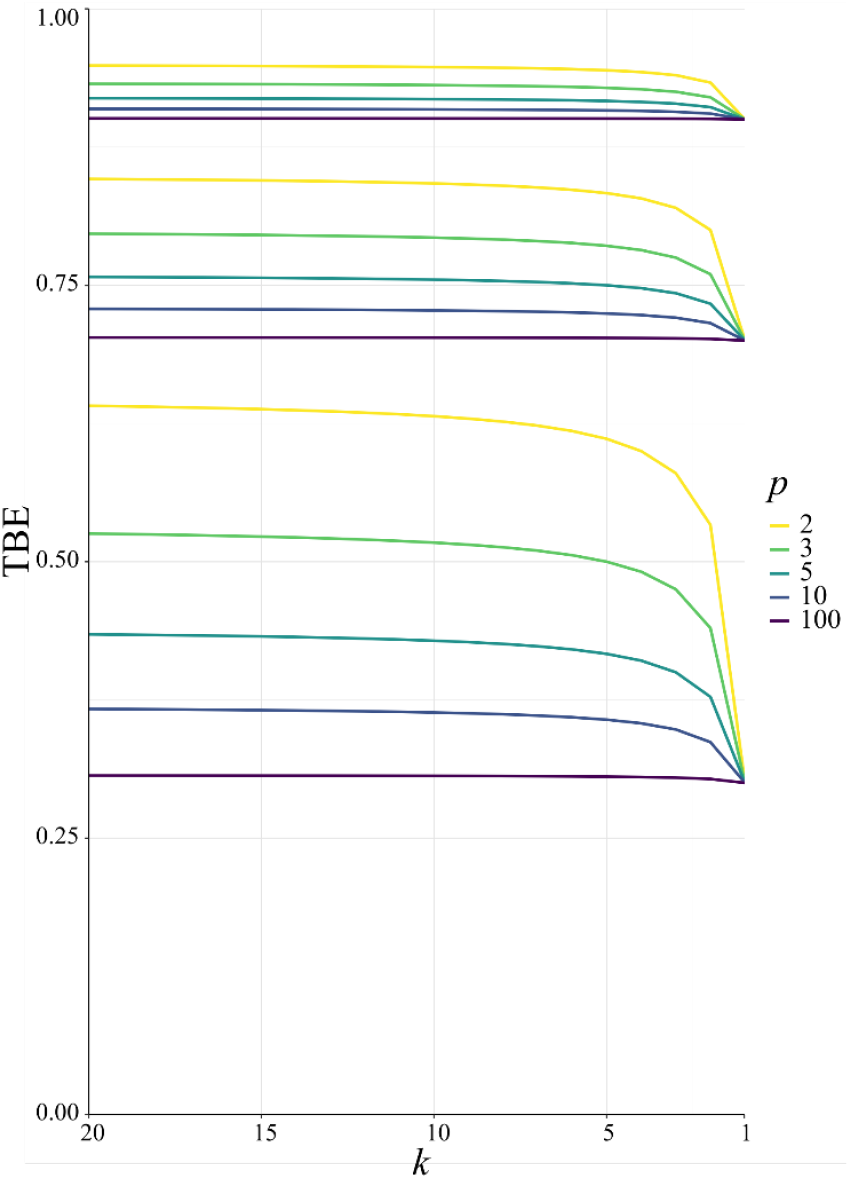
Variation of TBE supports in the presence of duplicates. Horizontal axis: *k* = 1 corresponds to the original tree with no duplicates, whereas with *k* = 20 each taxon is duplicated 20 times; the largest value of *k* is on the left to facilitate understanding of results with real data, where we progressively remove taxa (see below). Vertical axis: TBE support. Colorings: *p* corresponds to the depth of the branch in the original tree (e.g., *p* = 2 is a cherry, in yellow). The three sets of curves correspond to different values of the original TBE support without duplicates: *σ*(1) = 0.9 on top, 0.7 in the middle, 0.3 on bottom.

This simple model allows us to capture some trends in TBE supports. In line with Goldman’s hypothesis, the TBE supports of very small clades will indeed increase when duplicates are added to the tree, whereas in this model the FBP supports remain the same. However, this is much less true for medium-sized clades (e.g. 10-30 taxa), and for large clades the TBE supports are practically unchanged. Furthermore, the removal of duplicates before tree inference should be done systematically and is already considered good practice by most phylogeny software. We will confirm these results on TBE supports with our empirical data sets in the following sections. FBP is robust in this example, but not in our next example, which has no duplicates and a strong signal, but where we introduce a few rogue taxa.

### Presence of rogue taxa when the tree is globally supported

As in the previous example, let us consider again a bipartition P|Q, where P and Q have sampling size *p* and *q*, respectively, *p* ≤ *q* and *p* + *q* = *n*. We assume now that the signal is globally strong but affected by the presence of a few rogue taxa. Hence, the bipartition P|Q is ‘almost’ found in bootstrap trees, but every taxon has a small probability *π* (in the order of 1⁄*n*) of being rogue and randomly placed in P or Q in the bootstrap trees, following a uniform model with probability of being assigned to P = *p*⁄*n* and probability of being assigned to Q = *q*⁄*n*. In this model, we can easily compute FBP and TBE supports as a function of *π, p* and *n*.

If a taxon of P in the reference tree is rogue, its probability of being wrongly placed in a bootstrap tree is *q*⁄*n* = (*n* − *p*)⁄*n*. Conversely, the probability for a rogue taxon of Q to be wrongly placed is *p*⁄*n*. Thus, the total probability for a rogue to be wrongly placed is equal to 2p(*n* − *p*)⁄*n*^2^, and the probability for a taxon to be rogue and wrongly placed is equal to 2*πp*(*n* − *p*)⁄*n*^2^. Therefore, the expected FBP support is equal to the probability that no taxon is rogue and wrongly placed: FBP(*π, p, n*) = (1 − 2*πp*(*n* − *p*)⁄*n*^2^)^*n*^. As for TBE, the expected number of rogues is equal to *πn*, and so the TBE support is approximated by TBE(*π, p, n*) ≈ 1 − 2*π p*(*n* − *p*)/*n*(*p* − 1) (we assume here that *π* represents a small fraction of the taxa, making the perturbed-by-rogues version of P|Q the closest bipartition to P|Q in bootstrap trees).

Figure 2 shows the evolution of supports for FBP and TBE when varying *π* {0.001, 0.005, 0.01, 0.05} and *p* {2, 3 … 500} in a 1,000-taxon tree (*n* = 1000). With cherries (*p* = 2) both supports are the same, as explained before. For FBP, the support depends on the number of rogues (*π*), but also strongly on the sampling size *p* of the clade P. Figure 2 confirms that FBP drops extremely fast when the number of rogues is high (i.e., ∼50 rogues with *π* = 0.05). Furthermore, FBP also drops for large clades even with a very low number of rogues (e.g., ∼1 rogue with *π* = 0.001; then FBP < 70% when *p* > 232). The TBE support, on the other hand, behaves in a diametrically opposed way to the FBP support, but with much less variation; TBE rapidly increases when *p* is very low (between 2 and 5), and then slowly increases as *p* keeps increasing. Figure 2 highlights the very different nature of FBP and TBE supports in the presence of randomly distributed rogue taxa in large trees. TBE remains mostly stable no matter the depth of the reference branch (bipartition) when the number of rogue taxa is reasonable (i.e., 1 to 10 among 1000 taxa). On the contrary, the FBP support will always be considered low if there is even a single rogue when the topological depth of the reference branch is high (i.e., a large clade). These behaviors are clearly visible in our analyses of empirical datasets.

**Figure 2:**
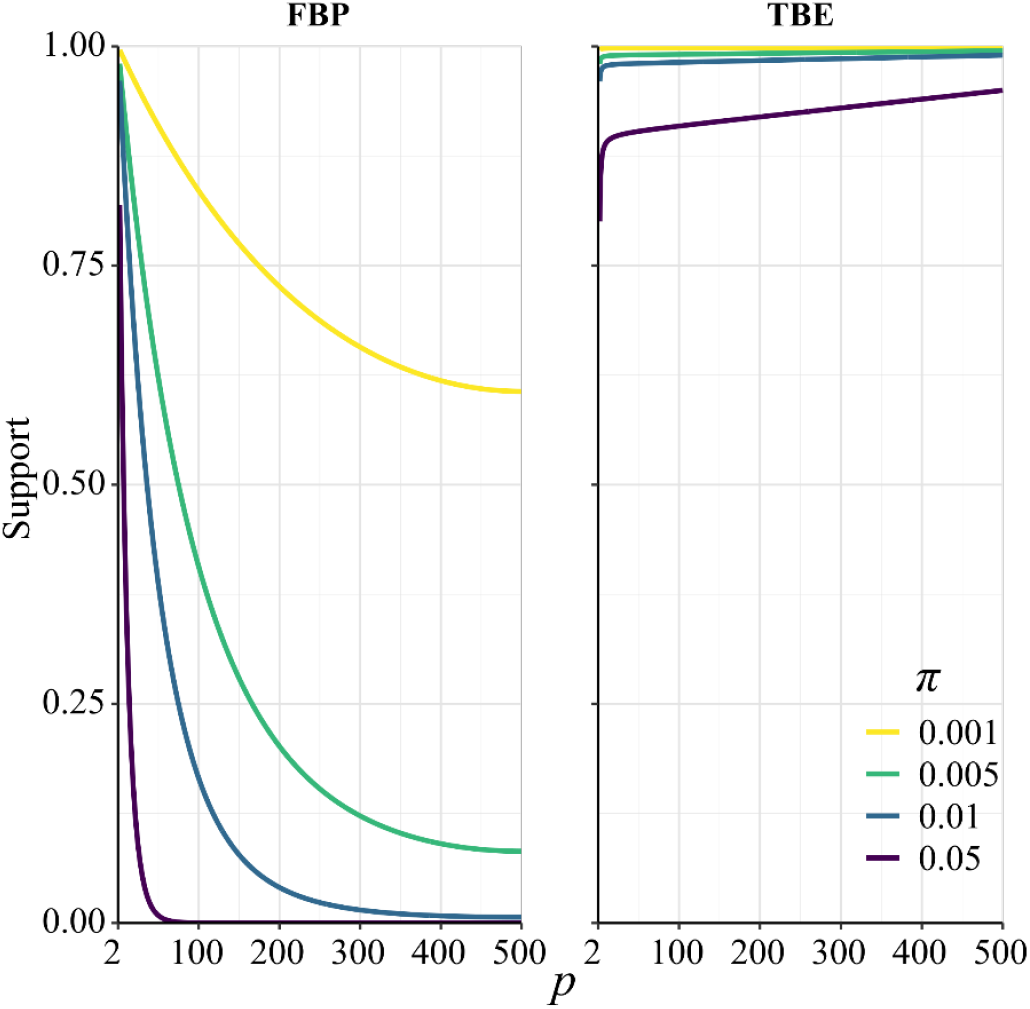
Theoretical results with strong signal but a few rogues. Evolution of support values (FBP on the left, TBE on the right) depending on sampling *p* of clade P, in a 1000-taxon tree. The color gradient represents the probability (*π*) of being rogue as the color darkens (e.g., *π* = 0.005 corresponds to ∼5 rogues among 1000 taxa).

In light of the two models presented above, there is reason to be concerned for both branch supports: for FBP, with deep branches and large clades in case of rogue taxa; for TBE, with shallow branches and small clades in case of sampling issues. However, real data are much more complex than these two theoretical examples, and thus we will further explore the properties of the two branch supports on a series of large biological datasets that supposedly have more or less sampling disequilibrium and rogue taxa.

## Results on empirical datasets

### Overview

Our dataset picking strategy consists in taking large empirical datasets from the literature, for which we expect some degree of heterogeneity in phylogenetic signal and sampling. This would imply that the genetic locus is expected to return good phylogenetic signal for some clades, but will fail to fully resolve a large portion of the phylogeny due to lack of sites. For example, barcode markers like *cytochrome c oxidase I* (COI) in most metazoans, or 16S ribosomal ribonucleic acid (16S rRNA) in prokaryotes, are typically adequate phylogenetic markers to get a rough idea of the relationships between taxa, but are not sufficient to build reliable phylogenies, especially with thousands of taxa. Moreover, unbalanced sampling is expected, with some clades being over-represented (e.g., Primates and Cetaceans in Mammals) and some others under-represented (e.g., Rodents).

We selected the two empirical datasets of the original TBE study (Lemoine et al. 2018) consisting of a COI mammal dataset (1,449 sequences) and an HIV dataset (9,147 *pol* sequences). Additionally, we used the full SARS-CoV-2 dataset from Zhukova et al. (2021) comprising 11,316 genomes. Finally, we selected 10 aligned nucleotide barcode datasets from Delsuc and Ranwez (2020) comprising between 1,000 and 2,000 mitochondrial COI sequences. All of these published datasets were already aligned and had in common to be large (> 1,000 sequences), with heterogeneous phylogenetic signal and unbalanced sampling.

Our methodology consisted in studying the evolution of FBP and TBE supports, starting from a reference tree with *N* taxa and unbalanced sampling, to reach a reduced and rebalanced target tree with *n* (<< *N*) taxa. In addition, along this path, we studied progressively more and more balanced intermediate trees. The sampling of these intermediate trees was defined by: *n*_*f*_ = *N* − *f* ∗ (*N* − *n*), with 0 ≤ *f* ≤ 1. For example, to get the midpoint between *N* and *n*, we choose *f* = 0.5, while *f* = 0 and *f* = 1 correspond to the starting and target trees, respectively. When we wanted to target the sampling of specific clades (e.g., HIV subtypes), then the representativeness of each clade was multiplied by *n* to obtain the required number *n*_*X*_ of sequences to be retained for clade *X*. Let *N*_*X*_ be the initial size of clade *X*, the intermediate clade samplings were then defined by *n*_*X,f*_ = *N*_*X*_ − *f* ∗ (*N*_*X*_ − *n*_*X*_). The target samplings (*n*_*X*_) in the reduced tree were defined by different criteria based on the specifics of the dataset: representativeness for mammals, worldwide prevalence for HIV subtypes, sampling density per country for SARS-CoV-2, and uniform sampling per delimited species for the barcode datasets.

The objective was to assess the stability and robustness of each branch-support measure throughout the subsampling. We selected a set of relevant clades for which one expects strong phylogenetic signal. For instance, long-branch clades like Cetacea or Marsupialia in the Mammals dataset are expected to be recovered even with a simple COI marker. The same applies to HIV subtypes and the *pol* gene. For such clades, with strong phylogenetic signal, a stable and robust branch support should give a high score, with more or less the same value, whatever the sampling size and balance.

To get supports for the target and intermediate trees, the ideal approach would be re-estimating the reference and bootstrap trees at each sampling point. However, this procedure can be quite computationally intensive, especially for very large datasets with multiple replicates. Another much faster approach is to prune the leaves of the starting reference and bootstrap trees, and calculate FBP/TBE supports on the reduced trees. The “pruned” approach is the one we favored to reduce computational costs, but in parallel, we also used the “re-estimated” approach on the target sampling to check if the results matched those obtained with the “pruned” approach. In the two approaches, all branches with length smaller than the expectation of having 0.5 mutations in total among all sites were collapsed in both the reference and bootstrap trees, to prevent adding noise in the support values (Guindon et al. 2010). The minimum branch length was defined by *ℓ* = 0.5⁄*L*, where *L* was the number of sites in the alignment. As such, *ℓ* is a simple form of confidence interval: branches with length less than *ℓ* are estimated to carry 0 mutations (i.e., no phylogenetic signal) and are collapsed, while branches with length greater than *ℓ* carry at least 1 mutation and are retained.

We looked for clades of interest in the reference trees (*T*) using well-established classifications. For mammals, we chose the NCBI taxonomy, with focus on Primates, Cetaceans, Marsupials, etc. For HIV, we used subtype annotation, and Nextstrain clades for SARS-CoV-2. These clades may (or not) be fully recovered in the reference tree, which is inferred from the data. To account for this, we traversed all branches in the reference tree and looked for the branch that minimized the transfer distance with the clade of interest (e.g., Primates with Mammals, or subtype B with HIV). We distinguished wrong taxa (i.e., taxa that do not belong to the clade of interest) from missing taxa (i.e., taxa that belong to the clade of interest, but are not found in the closest clade of *T*). The numbers of wrong and missing taxa are denoted *w* and *m*, respectively, and *w*+*m* is the transfer distance between the clade of interest and the closest clade in *T* that is selected. Hence, a clade in the reference tree that had no wrong and no missing taxa (*w* = *m* = 0) was perfectly recovered, with respect to the NCBI taxonomy, HIV subtypes, Nextrain clades, etc. Some other clades were not perfectly recovered in our reference trees, with a few wrong and missing taxa. In both cases, we measured the FBP and TBE supports of these reference clades, having in mind that a clade that is (almost) perfectly recovered should be highly supported. The same approach was used in Lemoine et al. (2018), for example with HIV, where several subtypes were not fully (but almost perfectly) recovered in the large tree estimated from the 9,147 available *pol* sequences (e.g., *w* = 2 and *m* = 0 with subtype B that comprises >3,500 sequences).

To summarize our overall strategy (Fig. 3), we begin with a large reference tree of sampling *N* (along with 1,000 bootstrap trees), and define an overall target sampling *n*. The reference and bootstrap trees of sampling *N* are then progressively pruned until achieving target sampling *n*, with multiple replicates to account for random effects. At each intermediate point, FBP/TBE values are computed from the pruned reference and bootstrap trees. Once target sampling is achieved, we re-estimate reference and bootstrap trees from sampling *n*, and compute again FBP/TBE values for comparison with the “pruned” approach.

**Figure 3:**
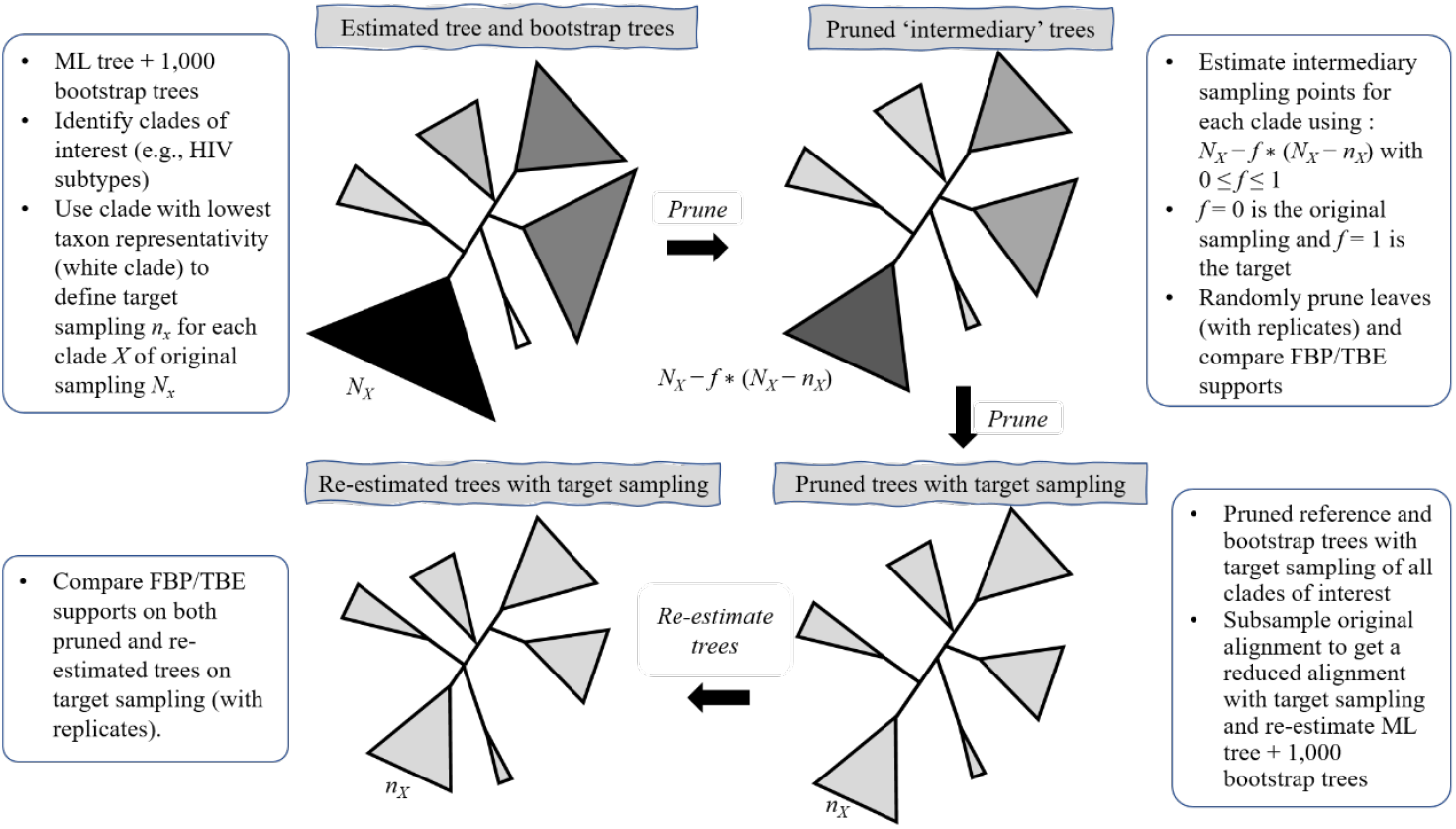
Summary of the overall sampling strategy. We start from a large reference tree of unbalanced sampling (*N*, top-left) and we define a target sampling (*n*). We then define intermediate sampling points (top-right) and prune the reference and bootstrap trees accordingly, across multiple replicates. When target sampling is achieved (bottom-right), we then subsample the original alignment accordingly and re-estimate a reference tree along with 1,000 bootstrap trees.

Prior to focusing on the selected clades, we briefly analyzed the overall average support in the Mammals, HIV and SARS-CoV-2 datasets, for different samplings and depths (Supp. Fig. SF1). Our results mainly show that FBP and TBE average supports remain stable (with a few exceptions) throughout the successive subsamplings (“pruned” experiments), even when the number of taxa is drastically reduced. Furthermore, we observe that the difference in average scores between the “pruned” and “re-estimated” experiments is low, thus validating our overall strategy (Fig. 3). Average TBE supports are on average higher than those of FBP, especially for deep branches (as expected, see above). Nevertheless, average FBP and TBE supports are low, even for shallow branches (<0.6 for FBP and <0.8 for TBE), as expected with these datasets. However, all these are average supports, and the results for the biologically important clades tell a very different story, as shown in the following.

### Mammals dataset

The Mammals dataset consists in 1,449 aligned COI protein sequences used in Lemoine et al. (2018). A few clades, almost without contradiction with the NCBI taxonomy (*w* ≈ *m* ≈ 0; for details, see Lemoine et al. 2018), had been highlighted to illustrate the differences between the FBP and TBE supports (Cetacea, Elephantidae, Geomyidae, “Insectivora”, Marsupialia, Monotremata, Mustelinae, Perognathinae and Simians). We selected these clades, but discarded those where the number of available species was insufficient for our experiments (i.e., Elephantidae and Monotremata). As expected, these clades have been unevenly sampled across the mammalian diversity, either over-sampled or under-sampled. For example, the Cetacea sampling represents 61% of all cetacean species, while the Soricomorpha (“Insectivora” in Lemoine et al. 2018) sampling represents only 13% of the Soricomorpha species.

We used the Integrated Taxonomic Information System (ITIS) to obtain the taxon representativeness *R*_*X*_ of each selected clade *X* (Supp. Tab. ST1). The target sampling *n* for the reduced tree from the initial sampling *N* (= 1,449) was estimated with the following steps. First, we selected the clade with the lowest sampling (i.e., Mustelinae, with *N*_*X*_ = 9). To obtain the smallest possible target sampling *n* while keeping all the selected clades, the target sampling *n*_*X*_ of Mustelinae was set to 2 (a cherry, the smallest possible clade). Then, we divided 2 by the Mustelinae taxon representativeness *R*_*X*_ = 0.0003 (i.e., 0.03% of Mammals diversity), to obtain the final target sampling of the reduced tree, *n* = 669. We estimated the target sampling of each clade *X* using *n*_*X*_ = *n* × *R*_*X*_. Finally, we estimated 3 intermediate samplings points with *f* = 0.25, 0.5 and 0.75. The species retained in the target sampling as well as in the intermediate points were randomly drawn from the initial species set, and this subsampling was iterated to obtain 30 replicates. This procedure allowed us to follow the trend of FBP and TBE scores while subsampling from initial unequal clade sampling (*N*_*X*_) to more accurate taxon representativeness (*n*_*X*_). FBP and TBE support values were computed at each intermediate point with the pruned approach (Fig. 3). We also re-estimated the reference tree and 1,000 bootstrap trees on the 30 replicates of the target sampling (669 taxa), and retrieved the selected clades in the reference tree as in (Lemoine et al 2018; see above and the Extended Material & Methods in the Appendix for details).

The initial results from Lemoine et al. (2018) showed that TBE significantly supports the selected clades while FBP does not, even when these clades were perfectly recovered in the reference tree (e.g., Cetacea, with *w* = *m* = 0). Our experiment confirms and complements these results by showing similar trends, when subsampling the initial species to achieve a (balanced) target sampling of the Mammal tree (Fig. 4). FBP tends to increase with reduced sampling, but in most cases it fails to reach the conventional threshold of 0.7 (e.g., the best score is Cetacea with original support ≈ 40% and average support with reduced sampling ≈ 60%). Two types of behavior can be observed for TBE, as predicted by our theoretical analyses (see above). When the number of taxa is sufficiently high (i.e., more than 10 in the reduced target sampling), TBE scores are high (>70%) and stable across all sampling points, as observed for the clades Cetacea, Marsupialia, Simians and Soricomorpha. However, when the number of taxa in the target sampling is low (<10), then TBE tends to decrease slightly and behave more like FBP as the number of taxa reduces (i.e., Geomyidae, Mustelinae, Perognathinae). When there are only 2 taxa left in the clade (i.e., Mustelinae), then FBP and TBE scores are equal, as expected and explained in the Introduction. Noteworthy, FBP shows more variability in maximum and minimum values (red ribbons) than TBE (green ribbons), even when there is no signal for the presence of rogues (*w* = *m* = 0, e.g., Cetacea and Geomyidae, Fig. 4). These results suggest a low robustness of FBP in the face of a heterogeneous phylogenetic signal, even when the clade is perfectly recovered in the reference tree. Lastly, the trees re-estimated at the target sampling (*n*) confirm our results obtained with the “pruned” approach, but tend to show greater variability in FBP scores across replicates, indicating that the robustness of FBP is even weaker in a realistic (full computation) setting with these mammalian data.

**Figure 4:**
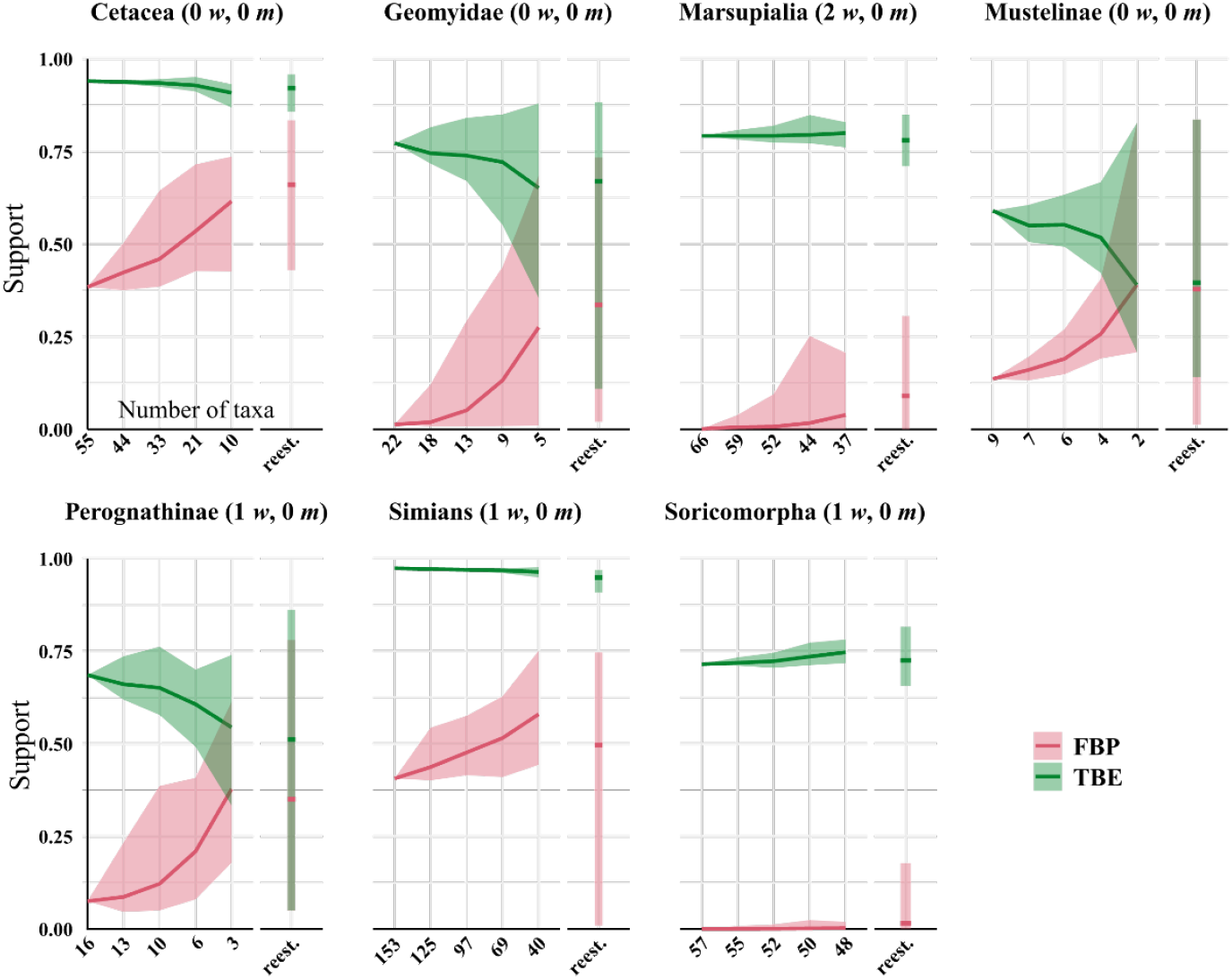
Results on the Mammals dataset. For each clade, the first value on the x-axis corresponds to the initial number of taxa and the last value to the number of taxa in the target sampling. We also provide the intermediate values, with proportions *f* = 0.25, 0.50 and 0.75. Trees were re-estimated on the target sampling (reest.). The number of wrong (*w*) and missing (*m*) taxa is indicated next to each clade name. To obtain the subsamples, taxa were randomly removed. The ribbon shows the maximum and minimum values over 30 replicates, the thick line is the mean value.

### The HIV dataset

Like the Mammals dataset, the HIV dataset was used in Lemoine et al. (2018) to highlight the differences between FBP and TBE. It consists of 9,147 HIV-1 group M *pol* sequences representing the 9 subtypes, and includes 50 recombinants detected using jpHMM (Schultz et al. 2009), that is, sequences that contain DNA from at least two different subtypes in the *pol* region. In this experiment, we achieved a representative sampling based on the current prevalence of each subtype in the worldwide population. The prevalence values of each subtype were obtained from Cassan et al. (2016, Fig. 3; Supp. Tab. ST2. Then, we applied the same approach as used with the Mammal dataset. We divided 2 by the prevalence of the most under-represented subtype (i.e., subtype K, with *R*_*X*_ = 0.1261%) to calculate *n* (= 1,599). We estimated 4 intermediate sampling points with *f* = 0.2, 0.4, 0.6 and 0.8, and achieved 30 replicates by randomly subsampling. With the target sampling, we re-estimated the trees and retrieved the selected clades as in (Lemoine et al. 2018; see above and the extended material & methods for details). In addition, we computed the average FBP/TBE support of all branches for different depths. This additional experiment allowed us to follow the evolution of the average FBP/TBE supports at different depths while reducing the sampling, instead of focusing only on HIV-1 group M subtypes that are known to be well supported in most datasets.

With the initial sampling, Lemoine et al. (2018) showed that all 9 subtypes were found in the reference tree and highly supported by TBE (although the average TBE support across the tree is relatively low; Supp. Fig. SF1). On the opposite, only the sparsely represented subtypes (i.e., F, H, J and K) were supported by FBP. A prominent example is the FBP/TBE supports of the B subtype, a clade comprising 3,559 sequences and which is almost perfect in the reference tree, since it contains all B sequences plus 2 B-recombinants (as detected by jpHMM). The FBP score of this clade is 0.03, indicating almost no signal, while the TBE score is 0.99, indicating a strong signal.

In the re-equilibrated trees using prevalence information, we see that the overall tendency observed in Lemoine et al. (2018) remains true, even when the sampling of some subtypes is considerably reduced (e.g., subtype B, from 3,559 to 226 sequences) and most recombinants are removed by random subsampling (Figure 5; Supp. Tab. ST2). In all cases, average TBE values are high (>90%) and with very little variation across replicates (green ribbons). In contrast, evolution of FBP supports across pruned trees does not follow the same pattern for all subtypes. In subtypes A, C, D or G, the FBP scores increase as sampling is reduced, but we observe a high variability (red ribbons) across replicates. Interestingly, those are the four subtypes (A, C, D, G) with a high number of recombinants (as detected with jpHMM; see Supp. Tab. ST3), with 4 to 12 wrong/missing taxa each (*w* + *m*; Fig. 5). Extreme variability across replicates is likely explained by the presence/absence of recombinants that are randomly subsampled. Quite differently, subtypes B and F (2 wrong in B corresponding to B-recombinants, none in F and no missing in both F and B) show little evolution of FBP support, and little variability across replicates, at least for the “pruned” strategy. This suggests that subtypes B and F might be affected by other unstable taxa than recombinants (or undetected recombinants), and numerous ones, so that scores across replicates are not affected by subsampling randomness. One could thus distinguish two forms of taxon instability (with the whole spectrum of intermediate cases): single rogue terminal taxa prone to instability (e.g., recombinants), and taxa belonging to globally unstable clades.

**Figure 5:**
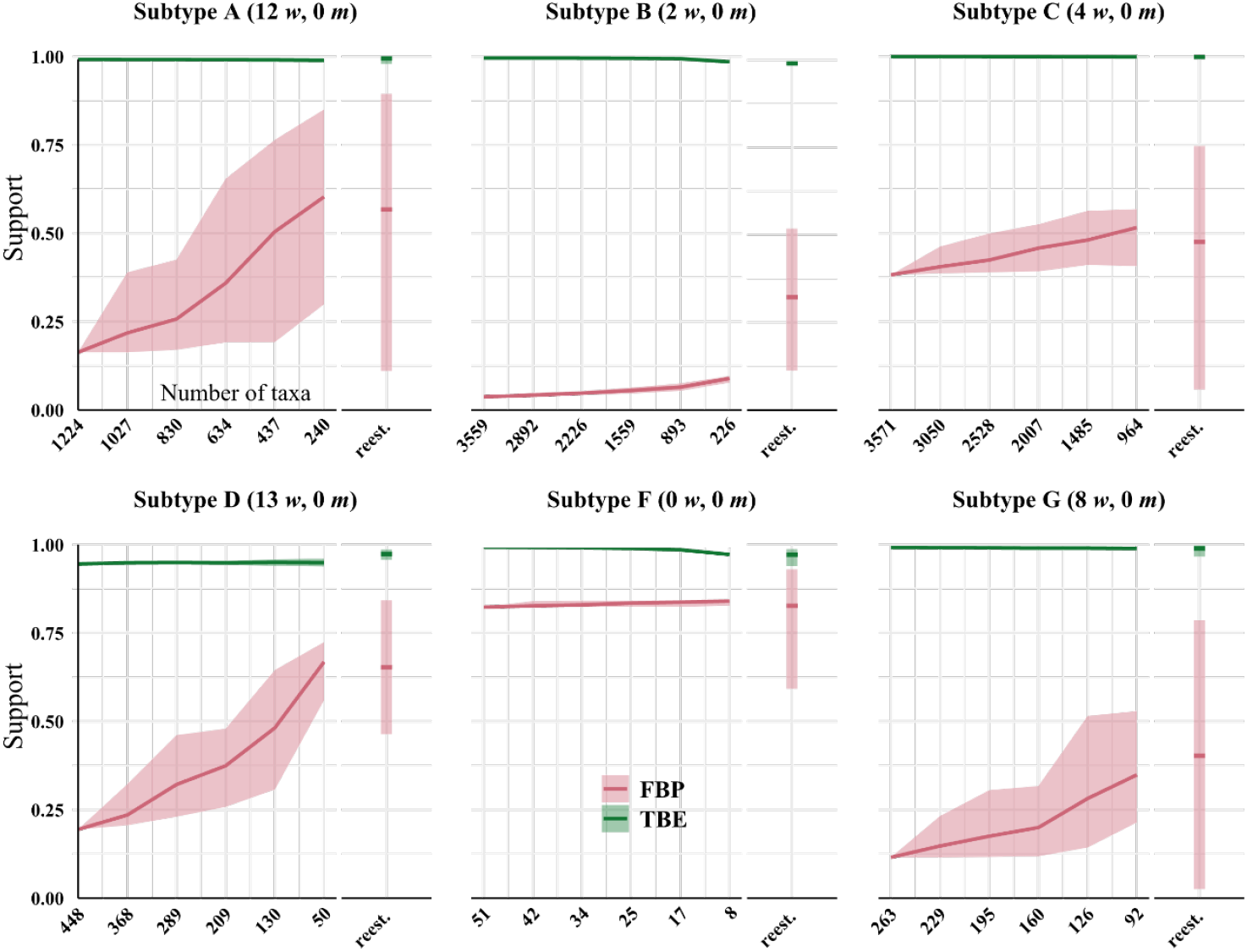
Results on the HIV dataset. We removed taxa within each HIV subtype based on prevalence. For each subtype (clade), the first value on the x-axis corresponds to the initial number of taxa and the last value to the number of taxa in the target sampling. We also provide intermediate values, corresponding to proportions *f* = 0.2, 0.4, 0.6 and 0.8 (see text for details). Trees were re-estimated on the target sampling (reest.). The number of wrong (*w*) and missing (*m*) taxa is indicated next to each subtype identifier. To obtain the subsamples, taxa were randomly removed. The ribbon shows the maximum and minimum values over 30 replicates, the thick line is the mean value.

Our results again illustrate the low robustness of FBP to rogue taxa and show how robust TBE is, whether or not some rogues are present (as predicted by our theoretical model with rogues, see above). One could argue that TBE might be over-supporting those clades, but our experiment on the overall support values across the tree show another story (Supp. Fig. SF1). The overall FBP/TBE support in the phylogeny is low for all ranges of branch depth (by the standards of each metric, i.e., less than 0.4 for FBP and less than 0.6 for TBE) and shows little change as trees are increasingly subsampled, except for cherries and deep branches where a trend emerges with this HIV dataset. Indeed, average FBP/TBE support for cherries (*p* = 2) slowly decreases as we remove more and more sequences. Conversely, average TBE supports for deep branches (*p* > 9) tend to increase. These overall trends contradict the results with selected clades/subtypes (Fig. 5), but are slight (less than 10% difference between initial and target samplings) and are not found in the analysis of the SARS-CoV-2 dataset that follows.

### The SARS-Cov-2 dataset

We retrieved the SARS-CoV-2 dataset from Zhukova et al. (2021) and processed the sequences through the Nextclade Web 2.3.0 interface (Aksamentov et al. (2021); https://clades.nextstrain.org) to assign each sequence to a Nextstrain “clade”. Some Nextstrain “clades” are paraphyletic, as they do not include the sequences of new child clades. When a “clade” was paraphyletic (e.g., clade 20B), we added all child clades in the sampling to make it monophyletic (i.e., 20D in the case of 20B). The result is a reference classification composed of 5 clades: 19B, 20A (= 20A+20B+20C+20D Nextstrain “clades”), 20C, 20B (= 20B+20D Nextstrain “clades”), and 20D. Sampling re-equilibrium had already been achieved by Zhukova et al. (2021), starting from the complete 11,316-sequence tree to 5 subsampled and re-balanced trees of size ∼2,000. The target sampling for these five datasets was defined to reflect the number of cases reported by country over time during the early months of the pandemic. We successively randomly pruned taxa by batches of ∼1,500 in the starting tree to retrieve the sampling of the 5 trees of size ∼2,000 from Zhukova et al. (2021), using 20 replicates each time, for a total of 20 ∗ 5 = 100 replicates. We extracted the FBP/TBE supports for the five selected clades (Fig. 6) and the average of these supports for different branch depth intervals (Supp. Fig. SF1), from the pruned and estimated trees. As with the other datasets, we retrieved the selected clades in the estimated trees by minimizing the number of wrong (*w*) and missing (*m*) taxa (see above and Appendix for details).

**Figure 6:**
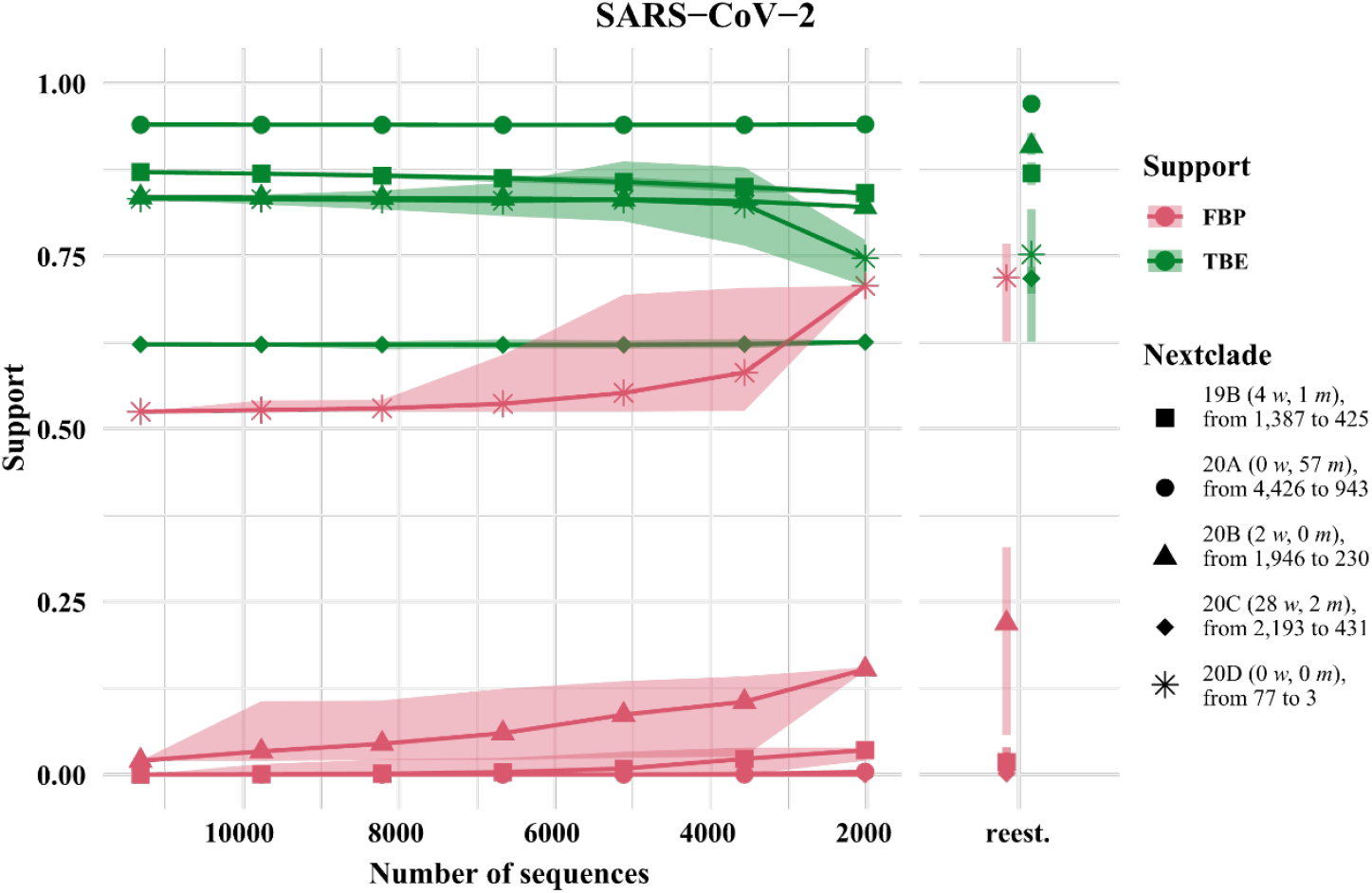
Results on the SARS-CoV-2 dataset. We randomly removed taxa from the original SARS-CoV-2 tree (11,316 tips) in order to achieve a better sampling equilibrium on trees with about 2,000 taxa. Trees were re-estimated on target sampling (reest.). The number of wrongly (*w*) and missing (*m*) taxa is given next to each clade identifier, and we provide the original and target samplings for each clade (e.g., ‘77 to 3’ with 20D). The ribbon indicates the maximum and minimum values, and the thick line corresponds to the mean value across all replicates.

The difference with the previously analyzed dataset (HIV) is that overall phylogenetic signal is scarce, with clades supported by only one or two mutations (https://nextstrain.org/; Supp. Tab. ST3). As for the previous datasets, most Nextstrain clades are well supported by TBE and not by FBP, even when the clade is well retrieved in the reference tree (e.g., 20D). Interestingly, support values are little affected by the number of wrongly placed and missing sequences (*w* + *m*), and the FBP/TBE scores remain stable across most clades upon subsampling (Fig. 6). For example, clade 20C is not well supported by TBE (∼0.62). Indeed, with 2,193 sequences in this clade, a score of 0.62 means that on average, (2193 – 1) x 0.38 = 833 sequences need to be transferred to recover that clade in the bootstrap trees; this number of 833 is much greater than the number of wrong/missing sequences in the reference clade (= 30, Fig. 6). As opposed to HIV, results with SARS-CoV-2 seem to indicate a global instability of a large number of taxa. This is likely due to the scarcity of the signal and the presence of “rogue clades”, which would be responsible for the transfer of hundreds of taxa in the bootstrap trees (thus greatly affecting TBE as well as FBP), rather than a few rogue terminal taxa as with HIV (e.g., subtype B, Fig. 5).

The evolution of average FBP/TBE support at different ranges of branch depth (Supp. Fig. SF1) indicates again that TBE is higher than FBP on average. FBP returns (again) heterogeneous average support values depending on the depth range (low FBP support for shallow branches, very low FBP support for deep branches), while TBE average supports are more homogeneous. However, these average results (Supp. Fig. SF1) confirm the finding in clade-specific Figure 6 that FBP and TBE supports are little affected by sampling biases. Moreover, both supports are higher in comparison with HIV (Supp. Fig. SF1), a difference which could be explained by the fact that SARS-CoV-2 trees on full genomes are easy to infer, even with parsimony (Thornlow et al. 2021), while HIV trees on a single marker might be more prone to saturation and other phylogenetic biases.

### The barcode datasets

We selected 10 aligned nucleotide barcode datasets from Delsuc and Ranwez (2020) comprising between 1,000 and 2,000 mitochondrial COI sequences (Acanthocephala, Archaeognatha, Bryozoa, Dermaptera, Megaloptera, Onychophora, Siphonaptera, Tardigrada, Testudines, Uraniidae). We estimated reference and 1,000 bootstrap trees on each datasets using RAxML-NG (Kozlov et al. 2019). We ran the maximum-likelihood heuristic of the multi-rate Poisson Tree Processes (mPTP) method (Kapli et al. (2017) in default mode to delimit putative species in each dataset. The output of mPTP was used to annotate species branches in each reference tree. Some of so delimited species have many sampled sequences (∼8% of retained species have >30 sequences), while others are poorly sampled (∼22% of retained species have 2 sequences; see also Supp. Fig. SF2). We computed FBP/TBE supports for each dataset on the “complete” trees with all sequences, and on the “deduplicated” trees were duplicated (i.e., strictly identical) sequences from the reference and bootstrap trees were removed. Our target sampling was defined by keeping a maximum of 5 different sequences (“max5”) for each species so delimited. The reference and bootstrap trees were fully re-estimated using RAxML-NG from this reduced and re-balanced set of sequences (the pruning approach was not used here). Sequences were not removed randomly, but using the greedy algorithm of (Steel 2005;(Pardi and Goldman 2005) implemented in the Phylogenetic Diversity Analyzer (PDA; Chernomor et al. 2015). This algorithm consists of iteratively deleting the taxon associated with the shortest branch; in doing so, we maximize the phylogenetic diversity of the tree while reducing the sampling. In total, our 10 barcode datasets add up to 15,111 COI sequences, of which 5,700 (∼38%) are duplicates, and a total of 1,390 putative species have been delineated. Some species clades were not retained in the analyses, because they were singletons in the complete or deduplicated trees, or because they were not found in the “max5” reference trees due to phylogeny re-estimation. This filtering downsized to 938 the number of putative species used in this experiment. We distinguished several categories of branches, based on the initial number of sequences in the delimited species, as well as supra-specific branches (i.e., branches above the species level).

On average, the FBP and TBE support values for the putative species branches are high, that is, above 0.7. This result was expected and is explained by our use of mPTP. This method counts the number of substitutions per branch and estimates the rates of branching events to detect which parts of the tree follow as speciation model (interspecific) and which follow a coalescent model (intraspecific). Thus, the delimited species branches usually correspond to multiple substitutions and carry a strong signal.

As expected, FBP scores do not change when removing duplicate sequences, since complete and deduplicated tree topologies are identical (Figure 7). We also notice that TBE scores only slightly decrease when removing duplicates (except for cherries). Thus, despite removing nearly 38% of the sequences, TBE shows support stability. The same stability in TBE scores is observed in the target sampling (“max5”) dataset, when reducing all species’ sampling to a maximum of 5 sequences. In contrast, FBP is much more affected than TBE by subsampling, especially for species that initially have many sequences (i.e., more than 30 samples). To ensure that observation, we computed the absolute delta difference between the initial and target (“max5”) sampling between all comparable branches across 10 datasets for each depth category. Results indicate a high delta in FBP scores between the initial and target sampling, particularly for species that are well represented in the initial sampling (with Δ = 0.19, versus 0.07 for TBE; Fig. 7).

**Figure 7:**
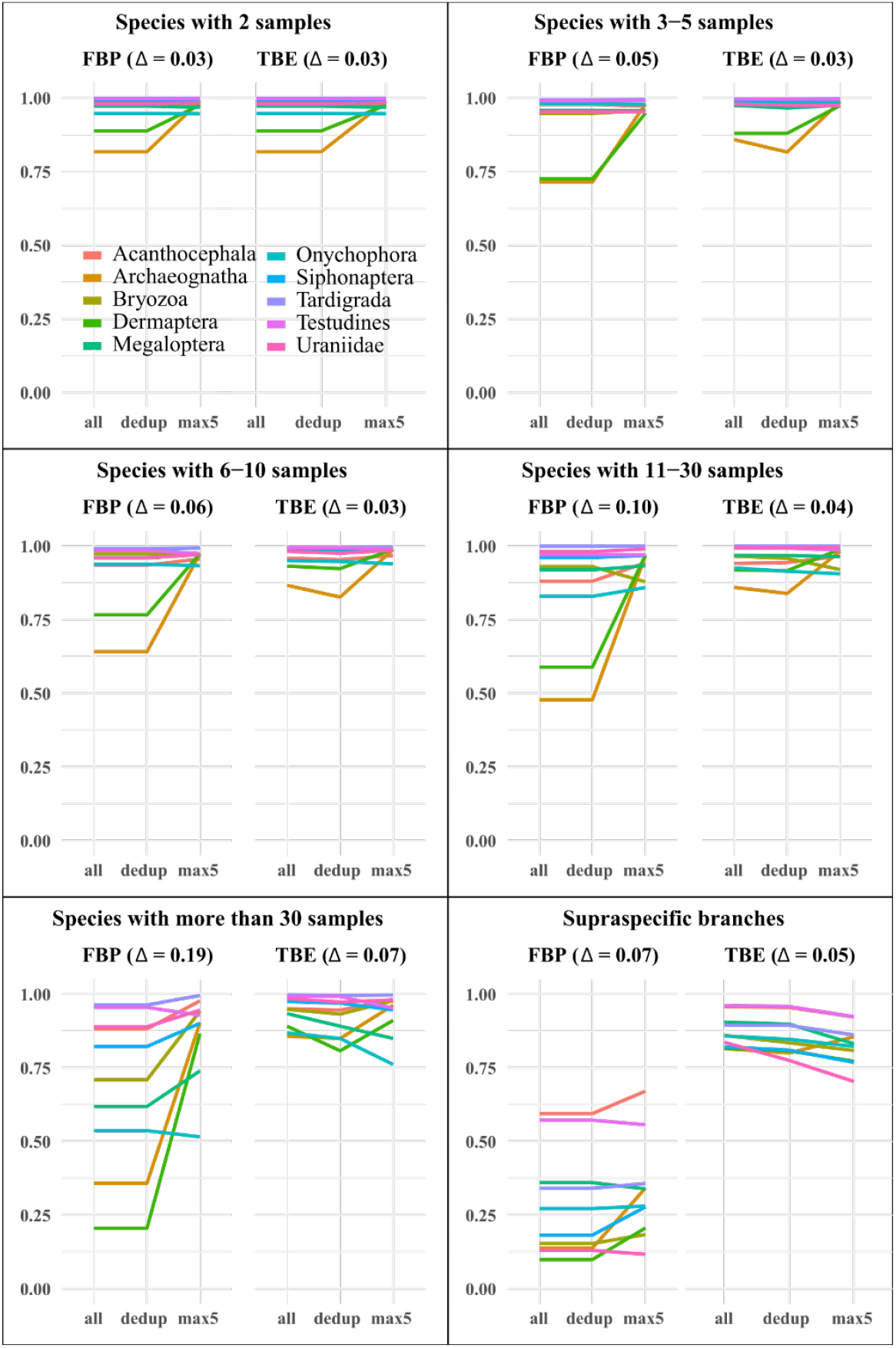
Results on the barcode datasets. Boxes correspond to the initial number of samples in the delimited species in the tree with all available sequences. E.g., if we consider the bottom-left panel “Species with more than 30 samples”, it means that for each tree on the full set of sequences (“all”), we retain all delimited species with more than 30 samples and report the mean support of the species branches on the first x-axis value. Second x-axis value takes the exact same set of species, but this time we compute supports using the trees with identical sequences removed (“dedup”). Finally, third x-axis value takes again the same set of species, but for re-estimated trees for which a maximum of 5 samples (“max5”) is kept by delimited species. “Supraspecific branches” are all branches above species level, i.e., we exclude species and infra-specific branches. Delta values correspond to the average absolute difference of supports between the initial (“all”) and target (“max5”) conditions, i.e., a high delta value indicates a large difference in FBP/TBE supports from a highly heterogeneously sampled dataset to a more homogeneous dataset.

This last experiment differs with the previous three in its subsampling strategy but also in terms of quantity of data and results. Instead of highlighting a few clades, we assume that most delimited species that are retrieved in all three modes of sampling should have a reasonably strong phylogenetic signal, should be well-supported (on average) and should be little affected by the subsampling procedure. While most species are well supported by both metrics, our results indicate a weak robustness of FBP to subsampling, particularly for species branches that initially contains many samples, while there is no reason to believe that these species in particular should be less supported than the others. This observation also holds for the supra-specific branches (Fig. 7), where TBE shows a better robustness (Δ = 0.05) than FBP (Δ = 0.07), even if the global tendency is consistent with the theoretical analyses, with TBE supports slightly decreasing with re-equilibrated (max5) sampling, while FBP increases by a rather larger margin in this condition.

## Discussion

Similar to what was shown in Lemoine et al. (2018), our theoretical and empirical results demonstrate the usefulness of TBE, especially for large trees with heterogeneous phylogenetic signal. Deep branches that are known to be essentially correct are generally supported by TBE, but FBP supports will generally be low, whether a phylogenetic signal is expected or not. In this study, we explored the impact of sampling biases on FBP/TBE support values. Through numerous datasets (1 Mammal, 1 HIV, 1 SARS-CoV-2 and 10 barcodes), we found no evidence that TBE falsely supports poor branches due to oversampling. In fact, most branches in these large datasets with heterogeneous and relatively low phylogenetic signal are on average poorly supported by TBE. Furthermore, our results indicate that TBE scores of the clades with significant signal are little affected by sampling biases, unless those clades are very small. Basically, in our experiments the TBE support remained unchanged when oversampled clades were downsampled to achieve balanced sampling, except for small clades whose TBE support tended to decrease. Indeed, for small clades the FBP and TBE supports are similar (they are identical for cherries). With medium-to-large clades, FBP supports were found to be not only lower but also much less robust than TBE supports, with an overall tendency to decrease with increasing sampling. Furthermore, depending on the (rogue) taxa sampled, the same clade with the same sample size can have low or high FBP support, revealing great variability in FBP support.

Based on these experiments, we suggest a few guidelines on how to conduct a routine bootstrap analysis to assess branch support in large trees:

- Remove duplicates: Duplicated (i.e., strictly identical sequences) are quite common in large datasets and can have an impact on TBE scores. Most ML software (e.g., IQ-TREE, RAxML-NG) already remove duplicate sequences prior to ML inference and then add back those sequences by creating near-zero length branches. We suggest that duplicates should also be removed from reference and bootstrap trees prior to computing TBE scores (and then added back).
- Collapse insignificant branches: In the obtained reference and bootstrap trees, many branches can be short and have no real biological meaning because they essentially correspond to no substitution event among all sites in the MSA.
- Systematically calculate both FBP and TBE: Bootstrap trees take a long time to compute, while calculating FBP/TBE is much faster in comparison, meaning that once bootstrap trees have been calculated, they should be used for both FBP and TBE. If the tree is highly supported by FBP (i.e., almost 100% for all branches), then TBE will not teach anything new. However, FBP should be systematically compared to TBE even with branches that appear to be “well” supported (e.g., 70%), because the interpretation of these two supports is very different. In this case, when TBE is close to 100%, this likely means that a few rogues are perturbing FBP analyses (as with HIV), whereas when TBE is also low (as with SARS-CoV-2), it is likely that many taxa are unstable and the overall phylogenetic signal is weak.
- Do not stop at the 70% threshold: For many phylogeneticists, 70% (or 80% for some) of FBP support has become the rule of thumb for assessing well-supported branches. This threshold with no statistical basis is likely inappropriate for many situations (signal or not, deep branches or not). For TBE, this value must be put in perspective with the sampling of the studied clade. With a clade of size *p* and of support *s*, one can easily calculate the average number of taxa that need to be transferred in the bootstrap trees to recover the reference clade, using the formula (*p* − 1) ∗ (1 − *s*). Thus, a score of 80% should not necessarily be considered as high support for TBE if the clade is very large, as this means that many taxa must be moved in the bootstrap trees to find the reference clade. These calculations are available in some software and websites (e.g., BOOSTER, https://booster.pasteur.fr/) and allow a better apprehension of TBE supports.

With now a better understanding of FBP and TBE behaviors under various sampling conditions, one of the major challenges in phylogenetics is to better interpret and use these branch supports. In the era of large-scale datasets, understanding the causes of a branch support to be low or high, through a better characterization of rogue taxa, would allow phylogeneticists to better comprehend their data and their flaws.

## Data availability

All data and scripts will be available upon publication on Dryad (https://doi.org/10.5061/dryad.ncjsxkt05), and can be requested from the authors.

## Appendix

Extended Material & Methods, Supplementary Figures (SF1, SF2) and Tables (ST1, ST2, ST3) are available in the Appendix.

## Acknowledgments

Computations have been performed on the Plateforme de Calcul Intensif & Algorithmique (PCIA, UAR 2700 2AD – Muséum National d’Histoire Naturelle, Centre National de la Recherche Scientifique, Paris, FRANCE) and on the cluster of the Institut Pasteur (Paris, France).

## Appendix

### Extended Material & Methods

**[*additional information about the M&M are added in italic*]**

#### The Mammals dataset

The Mammals dataset consists in 1,449 aligned COI protein sequences used in Lemoine et al. (2018). A few clades, almost without contradiction with the NCBI taxonomy (*w* ≈ m ≈ 0; for details, see Lemoine et al. 2018), had been highlighted to illustrate the differences between the FBP and TBE supports (Cetacea, Elephantidae, Geomyidae, “Insectivora”, Marsupialia, Monotremata, Mustelinae, Perognathinae and Simians). We selected these clades, but discarded those where the number of available species was insufficient for our experiments (i.e., Elephantidae and Monotremata). As expected, these clades have been unevenly sampled across the mammalian diversity, either over-sampled or under-sampled. For example, the Cetacea sampling represents 61% of all cetacean species, while the Soricomorpha (“Insectivora” in Lemoine et al. 2018) sampling represents only 13% of the Soricomorpha species.

We used the Integrated Taxonomic Information System (ITIS) to estimate the taxon representativeness *R*_*X*_ of each selected clade *X* (see Table ST1; *Retrieved June, 5*^*th*^, *2022, from the ITIS*, *www.itis.gov*, *CC0* https://doi.org/10.5066/F7KH0KBK). The target sampling *n* for the reduced tree from the initial sampling *N* (= 1,449) was estimated with the following steps. First, we selected the clade with the lowest sampling (i.e., Mustelinae, with *N*_*X*_ = 9). To obtain the smallest possible target sampling *n* while keeping all the selected clades, the target sampling *n*_*X*_ of Mustelinae was set to 2 (a cherry, the smallest possible clade). Then, we divided 2 by the Mustelinae taxon representativeness *R*_*X*_ = 0.0003 (i.e., 0.03% of Mammals diversity), to obtain the final target sampling of the reduced tree, *n* = 669. We estimated the target sampling of each clade *X* using *n*_*X*_ = *n* × *R*_*X*_. Finally, we estimated 3 intermediate samplings points with *f* = 0.25, 0.5 and 0.75 *(while f = 0 and f = 1 correspond to the starting and target trees, respectively)*. The species retained in the target sampling as well as in the intermediate points were randomly drawn from the initial species set, and this subsampling was iterated to obtain 30 replicates. This procedure allowed us to follow the evolution of FBP and TBE scores while subsampling from initial unequal clade sampling (*N*_*X*_) to more accurate taxon representativeness (*n*_*X*_).

*In Lemoine et al. (2018), bootstrap trees were initially estimated with RAxML version 8 (Stamatakis 2014) but using rapid bootstrapping (Stamatakis et al. 2008) and thus bootstrap trees were lacking branch lengths. We kept the initial reference tree from Lemoine et al. (2018) but re-estimated 1,000 ‘traditional’ bootstrap trees using the following commands:*

1. *Building the bootstrap alignments with goalign 0*.*3*.*6a* goalign build seqboot -i align.fasta -n 1000 -o align_boot_
2. *Inferring the reference and bootstrap trees with RAxML 8*.*2*.*8*

raxmlHPC-PTHREADS-AVX2 -m PROTGAMMAWAG -c 6 -s align.fasta -n NAME -T 4 -p 123456789

*Reference and bootstrap trees were also re-estimated on alignments with target sampling using the same RAxML command. Trees were pruned using “gotree prune” (see “List of Commands” section). Prior to each FBP and TBE support computation, branches shorter than 0*.*000949 (=0*.*5/527) were collapsed using “gotree collapse length” (see “List of Commands” section). This value corresponds to the expectation of having less that 0*.*5 mutations on the branch, given that the alignment has 527 sites*. FBP and TBE support values were computed at each intermediary point with the pruned approach (Fig. 3) *using option “gotree compute support fbp/tbe” (see “List of Commands” section)*.

We also re-estimated reference and 1,000 bootstrap trees on the 30 replicates of the target sampling (669 taxa) *using the same RAxML command as above and* retrieved the selected clades in the reference tree as in Lemoine et al (2018). *To retrieve clades that are closest to the reference NCBI taxonomy, we used “gotree compare edges” (see “List of Commands” section) which looks for the edge in the compared tree that minimizes the transfer distance with the reference clade in the NCBI taxonomy. To achieve that, trees must have the same set of leaves, so for reduced trees we pruned the NCBI taxonomy accordingly using “gotree prune”. The “gotree compare edges” command outputs taxa to transfer for each edge with either a plus sign “+” or minus sign “-”. A “+” sign indicates that this taxon should be added to the compared edge in order to retrieve the reference edge, we call these taxa missing taxa (m), i*.*e*., *taxa that belong to the reference clade but are not found in the closest edge of the estimated tree. A “-” sign indicates that a taxon found in the compared tree should be removed in order to retrieve the reference clade, we called these wrong taxa (w), i*.*e*., *taxa that do not belong to the reference clade*.

#### The HIV dataset

Like the Mammals dataset, the HIV dataset was used in Lemoine et al. (2018) to highlight the differences between FBP and TBE. It consists of 9,147 HIV-1 group M *pol* sequences representing the 9 subtypes, and includes 50 recombinants detected using jpHMM (Schultz et al. 2009), that is, sequences that contain DNA from at least two different subtypes in the *pol* region. In this experiment, we achieved a representative sampling based on the current prevalence of each subtype in the worldwide population. The prevalence values of each subtype were obtained from Cassan et al. (2016, Fig. 3; Table ST2). Then, we applied the same approach as used with the Mammal dataset. We divided 2 by the prevalence of the most under-represented subtype (i.e., subtype K, with *R*_*X*_ = 0.1261%) to calculate *n* (= 1,599). We estimated 4 intermediate sampling points with *f* = 0.2, 0.4, 0.6 and 0.8, and achieved 30 replicates by random subsampling. With the target sampling, we re-estimated the trees and retrieved the selected clades as in (Lemoine et al. 2018; see Table ST1 for details). *For the HIV dataset, bootstrap trees were initially estimated with FastTree version 2*.*1*.*11 (Price et al. 2010) without using the double precision algorithm. By default, standard FastTree does not allow branch lengths of less than 0*.*0005 — corresponding to the expectation of having 0*.*52 mutations on that branch, given that the alignment has 1,043 sites. Since our rule of thumb is to collapse branch lengths with expectation of having less than 0*.*5 mutations, we re-estimated 1,000 bootstrap trees on the 9,147 HIV-1 M pol sequences using FastTree version 2*.*1*.*11 Double precision:*

~~~
FastTreeDbl -nopr -nosupport -gtr -nt -gamma <bootstrap alignment>
~~~

*Reference and bootstrap trees were also re-estimated on alignments with target sampling using the same command. To maximize branch length estimation precision, we re-estimated branch lengths on the FastTree topology using RAxML-NG v. 1*.*1*.*0 (Kozlov et al. 2019) on all (re)estimated and pruned trees:*

~~~
raxml-ng --evaluate --msa <alignment> --tree <FastTree tree> --model GTR+G
~~~

*As for the Mammals dataset, trees were pruned using “gotree prune” and compared to a reference taxonomy of the 9 subtypes using “gotree compare edges”*. In addition, we computed the average FBP/TBE support of all branches for different depths. *Support values for each edge and depth values were computed using “gotree stats edges” (see “List of Commands” section)*. This additional experiment allowed us to follow the evolution of the average FBP/TBE supports at different depths while reducing the sampling, instead of focusing only on HIV-1 group M subtypes that are known to be well supported in most datasets.

#### The SARS-CoV-2 dataset

We retrieved the SARS-CoV-2 dataset from Zhukova et al. (2021) and processed the sequences through the Nextclade Web 2.3.0 interface (Aksamentov et al. (2021); https://clades.nextstrain.org) to assign each sequence to a Nextstrain “clade”. Some Nextstrain “clades” are paraphyletic, as they do not include the sequences of new child clades. When a “clade” was paraphyletic (e.g., clade 20B), we added all child clades in the sampling to make it monophyletic (i.e., 20D in the case of 20B). The result is a reference classification composed of 5 clades: 19B, 20A (= 20A+20B+20C+20D Nextstrain “clades”), 20C, 20B (= 20B+20D Nextstrain “clades”), and 20D. Sampling re-equilibrium had already been achieved by Zhukova et al. (2021), starting from the complete 11,316-sequence tree to 5 subsampled and re-balanced trees of size ∼2,000. The target sampling for these five datasets was defined to reflect the number of cases reported by country over time during the early months of the pandemic.

*For the SARS-CoV-2 dataset we ran multiple tests to determine what would be the best approach to infer the tree on the 8,541 genomes — after removing duplicate genomes from the initial 11,316 genomes dataset of Zhukova et al. (2021). We looked at the best trade-off for running time and tree accuracy between FastTree, IQ-TREE 2 (Minh et al. 2020), IQ-TREE 2 in fast mode and RAxML-NG (results not shown). Tree accuracy was evaluated by counting the number of taxa to transfer from the estimated tree to recover the Nextstrain clades from our reference taxonomy. The fastest approaches were by far FastTree and IQ-TREE 2 -fast, but the total number of taxa to transfer for FastTree (309) was much higher than for IQ-TREE (82). The total number of taxa to transfer was not much different for other, more time consuming, methods, hence IQ-TREE 2 -fast was selected as the main approach for the SARS-CoV-2. ModelFinder (implemented in IQ-TREE 2) was used to select the best substitution model on the overall alignment:*

~~~
iqtree2 -nt AUTO -s <alignment> -m MF
~~~

*The substitution model GTR+F+I+R3 was retained best using both Akaike and bayesian information criterion. We then launched tree inference on reference and bootstrap trees using:*

~~~
iqtree2 -fast -nt AUTO -s <alignment> -m GTR+F+I+R3
~~~

We then successively randomly pruned tips by batches of ∼1,500 in the starting tree *using “gotree prune”* to retrieve the sampling of the 5 trees of size ∼2,000 from Zhukova et al. (2021), using 20 replicates each time, for a total of 20 ∗ 5 = 100 replicates. We extracted average FBP/TBE overall supports from the pruned and re-estimated trees (see Figure SF1). *Support values for each edge and depth values were computed using “gotree stats edges”*.

#### The barcode datasets

We selected 10 aligned nucleotide barcode datasets from Delsuc and Ranwez (2020) comprising between 1,000 and 2,000 mitochondrial COI sequences (Acanthocephala, Archaeognatha, Bryozoa, Dermaptera, Megaloptera, Onychophora, Siphonaptera, Tardigrada, Testudines, Uraniidae). *For the 10 barcode datasets, we first removed duplicated (i*.*e*., *strictly identical) sequences from the initial alignment using “goalign dedup” (see “List of Commands” section)*. We estimated reference and 1,000 bootstrap trees on each datasets using RAxML-NG (Kozlov et al. 2019) :

~~~
raxml-ng --all --thread 16 --msa <alignment> --model GTR+G --bs-trees 1000
~~~

*Trees were then ‘repopulated’ using “gotree repopulate” (see “List of Commands” section) to add back the duplicated sequences*. We ran the maximum-likelihood heuristic of the multi-rate Poisson Tree Processes (mPTP) method (Kapli et al. (2017) in default mode to delimit putative species in each dataset :

~~~
mptp --ml --multi --tree_file <tree>
~~~

The output of the mPTP run was used to annotate species edges in each reference tree. *We used “gotree annotate” (see “List of Commands” section) to annotate each edge corresponding to a delimited species and thus distinguish species edges from intra- and supra-specific edges*. Some of so delimited species have many sampled sequences (∼8% of retained species have >30 sequences), while others are poorly sampled (∼22% of retained species have 2 sequences; see Figure SF2). We computed FBP/TBE supports for each dataset on the “complete” trees with all sequences, and on “deduplicated” trees were duplicated (i.e., strictly identical) sequences from the reference and bootstrap trees were removed. Our target sampling was defined by keeping a maximum of 5 different sequences (“max5”) for each species so delimited. The reference and bootstrap trees were fully re-estimated using RAxML-NG from this reduced and re-balanced set of sequences (the pruning approach was not used here). Sequences were not removed randomly, but using the greedy algorithm of (Pardi and Goldman 2005; Steel 2005) implemented in the Phylogenetic Diversity Analyzer (PDA; Chernomor et al. 2015). This algorithm consists of iteratively deleting the taxon associated with the shortest branch; in doing so, we maximize the phylogenetic diversity of the tree while reducing the sampling. *For species edges comprising more than 5 taxa, we extracted the corresponding subtrees using “gotree prune” and ran PDA to reduce the number of samples per species to 5 while maximizing diversity:*

~~~
pda -g -k 5 <species level subtree>
~~~

*We re-estimated trees using the same RAxML-NG command as above on trees where a maximum of 5 samples per species were kept. Finally, we compared species edges between three alternative samplings (“full”, “dedup” and “max5”) and considered only the edges that were found in all three samplings. To get support information about supra-specific edges, i*.*e*., *edges above the species level, we randomly selected 1 sample per species in each “full”, “dedup” and “max5” datasets using “gotree prune”, and then extracted support information for all internal edges using “gotree stats edges”. We also computed the absolute delta difference between the “full” and “max5” samplings by first reducing the “full” tree sampling to the “max5” target sampling (using “gotree prune”), then we compared the two trees using “gotree compare edges”, and finally we summarized the support values of all comparable edges across the 10 datasets for each depth category (2, 3-5, 6-10, 11-30, >30, supraspecific)*.

#### List of commands

For tree manipulation and description, we mainly used Gotree/Goalign (Lemoine and Gascuel 2021), a toolkit specifically designed for phylogenetic workflows.

Deduplicate sequences that present identical sequences, i.e., identical sequences ID are indexed in a log file.

~~~
goalign dedup -i <alignment> -l <output log>
~~~

After tree estimation, sequences can then be added back to the tree using Gotree :

~~~
gotree repopulate -i <tree> -g <file with identical tips>
~~~

Take a subset of sequences from the input alignment using a list of terminal taxa:

~~~
goalign subset -i <alignment> -f <list of sequences>
~~~

To get 1,000 bootstrap alignments from an input alignment:

~~~
goalign build seqboot -i <alignment> -n 1000 -S
~~~

A minimum branch length threshold is calculated for each tree; if a branch length is below that threshold then the branch is collapsed. The minimum length *ℓ* is determined by the following formula: *ℓ* = 0.5⁄*L*, where L is the number of sites in the alignment. To collapse branch length, we use:

~~~
gotree collapse length -i <tree> -l <*ℓ*>
~~~

Compute FBP and TBE supports from reference and bootstrap trees:

~~~
gotree compute support fbp -t 10 -i <reference tree> -b <bootstrap trees>
gotree compute support tbe -t 10 -i <reference tree> -b <bootstrap trees>
~~~

Prune a list of terminal taxa from an input tree:

~~~
gotree prune -i <tree> -f <list of tips>
~~~

To annotate a branch (edge) in a rooted tree, we need a file with one line per internal branch (light side) to annotate with a corresponding list of tip names. Gotree finds the LCA of every tips whose name is in the list:

~~~
gotree annotate -i <tree> -m <map file for annotation>
~~~

To select a subtree from a (rooted) input tree whose root has a given name:

~~~
gotree subtree -i <tree> -n <name of edge>
~~~

Extract statistics such as edge length, support, name, or depth (number of tips on the lightest side of a split) for each edge in a tree:

~~~
gotree stats edges -i <tree>
~~~

Compare all branches (edges) from a tree against a reference tree:

~~~
gotree compare edges -i <reference tree> -c <compared tree>
~~~

**Figure SF1:**
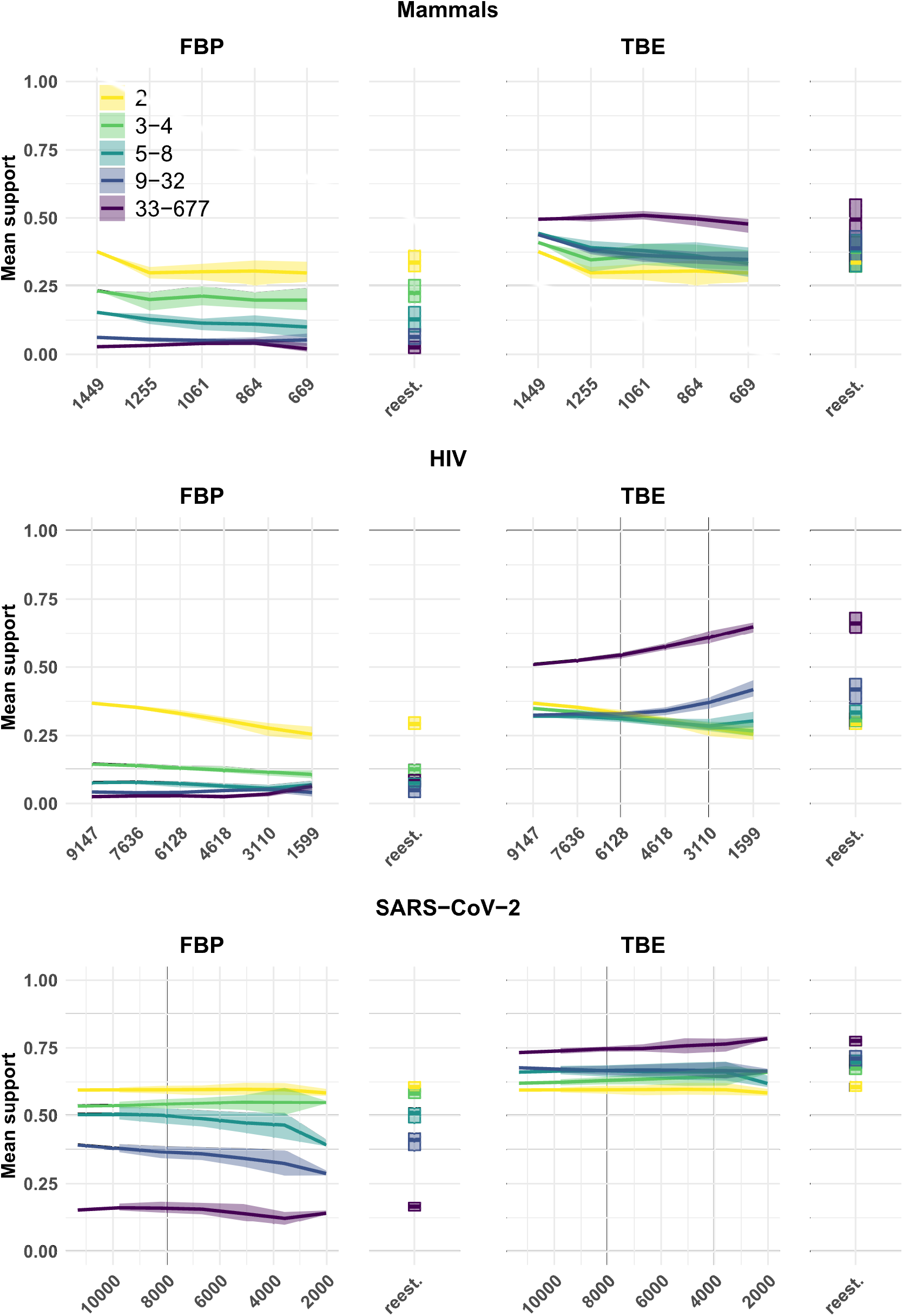
Overall global support values. X-axes correspond to the total number of leaves in the tree. Colors indicate the number of taxa on the light side of a branch. The ribbon indicates the maximum and minimum values across all replicates, and the thick line corresponds to the mean value across all replicates.”reest.” corresponds to the values on the reestimated trees on target sampling.

**Table ST1:** Taxon representativeness for highlighted clades, original and target sampling. nb-species: total number of Mammals species in the clade; %Representativeness: percentage of the number of species in the clade divided by the total number of Mammals (ITIS – 05/06/2022); nb-seq: number of sequences in the clade that belongs to the starting reference tree; %nb-seq/nb_species: percentage of the number species in the clade, represented by sequences in the starting reference tree (e.g. 61% of Cetacea are present, while only 13% of Soricomorpha are represented); Wrong (*w*) and Missing (*m*) are obtained by comparing the starting reference tree with the NCBI taxonomy (e.g., the Cetacea clade is perfectly recovered, *w* = *m* = 0, while 2 taxa have to be removed from the best clade in the starting reference tree to recover the Marsupiala; see extended M&M for details); Total: size (N_X_) of the best clade in the starting reference tree (e.g., 64+2 for the Marsupiala); Target: number (n_X_) of sequences of the clade in the target sampling, that is equal to 11-12% of the species diversity in the clade; %n_x_ / N_X_: percentage of the number of sequences kept in the target sampling to have nearly the same diversity in all clades (e.g., only 18% of the Cetacea are kept, while we kept 84% of the Soricomorpha). We see here how the target sampling re-balances the sampling across the clades of interest. In the target sampling, every clade has a number of sequences representing approximatively 11-12% of its species diversity.

| Clades | nb_species | %Representativeness | nb_seq | %nb_seq / nb_species | Wrong | Missing | Total (N<sub>x</sub>) | Target (n<sub>x</sub>) | %n<sub>x</sub> / N<sub>x</sub> |
| --- | --- | --- | --- | --- | --- | --- | --- | --- | --- |
| Cetacea | 90 | 14.9 | 55 | 61 | 0 | 0 | 55 | 10 | 18 |
| Geomyidae | 41 | 6.8 | 22 | 53 | 0 | 0 | 22 | 5 | 22 |
| Marsupialia | 335 | 5.6 | 64 | 19 | 2 | 0 | 66 | 37 | 56 |
| Mustelinae | 18 | 0.30 | 9 | 50 | 0 | 0 | 9 | 2 | 22 |
| Perognathinae | 27 | 0.45 | 15 | 55 | 1 | 0 | 16 | 3 | 18 |
| Simians | 363 | 6.0 | 152 | 41 | 1 | 0 | 153 | 40 | 26 |
| Soricomorpha | 428 | 7.1 | 56 | 13 | 1 | 0 | 57 | 48 | 84 |
| Remaining | 4723 | 78.4 | 1076 | 22 |  |  | 1071 | 525 | 48 |
| Total | 6025 |  | 1449 |  |  |  | 1449 | 669 | 46 |

**Table ST2:** HIV sampling, prevalence by subtype and target sampling. jpHMM: Sequences were assigned to subtypes (or else classified as recombinants) using jpHMM (see Lemoine et al. 2018 for further details); Prevalence (PNAS): worldwide prevalence of each subtype as extracted from Cassan et al. (2016; Fig. 3); Prevalence (rescaled): rescaled prevalence values for the purpose of our experiment; Wrong (*w*) and Missing (*m*) are obtained by comparing the starting reference tree with the jpHMM classification; Total: size (N_X_) of the best clade in the starting reference tree (e.g., 1212+12 for the subtype A), with outliers corresponding to wrong sequences; Target: number (n_X_) of sequences of the subtype clade in the target sampling, which is equal to the rescaled prevalence of the subtype (e.g., 15% for subtype A) multiplied by the total number of sequences in the target (1599, see text for details); %n_x_ / N_X_: percentage of the number of sequences kept in the target sampling to have nearly the correct prevalence in all clades (e.g., only 6% of the subtype B sequences are kept, while we kept 35% subtype G sequences). We see here how the target sampling re-balances the sampling across the clades of interest based on prevalence: some overrepresented subtypes (e.g., subtype B) have a considerably reduced sampling compared to other subtypes (e.g., subtypes K, J, G, C or A).

| Subtype | jpHMM | Prevalence (PNAS) | Prevalence (rescaled) | Wrong | Missing | Total (N<sub>x</sub>) | Target (n<sub>x</sub>) | %n<sub>x</sub> / N<sub>x</sub> |
| --- | --- | --- | --- | --- | --- | --- | --- | --- |
| A | 1212 | 0.12 | 0.151 | 12 | 0 | 1224 | 240 | 19.6 |
| B | 3557 | 0.113 | 0.142 | 2 | 0 | 3559 | 226 | 6.4 |
| C | 3567 | 0.482 | 0.608 | 4 | 0 | 3571 | 964 | 27.0 |
| D | 435 | 0.025 | 0.032 | 13 | 0 | 448 | 50 | 11.2 |
| F | 51 | 0.004 | 0.005 | 0 | 0 | 51 | 8 | 15.7 |
| G | 255 | 0.046 | 0.058 | 8 | 0 | 263 | 92 | 35.0 |
| H | 12 | 0.001 | 0.001 | 0 | 2 | 10 | 2 | 20.0 |
| J | 6 | 0.001 | 0.001 | 0 | 0 | 6 | 2 | 33.3 |
| K | 2 | 0.001 | 0.001 | 0 | 0 | 2 | 2 | 100.0 |
| Recombinants | 50 |  |  |  |  |  |  |  |
| Outliers |  |  |  |  |  | 13 | 13 |  |
| TOTAL | 9147 |  |  |  |  | 9147 | 1599 | 17.5 |

**Table ST3:**
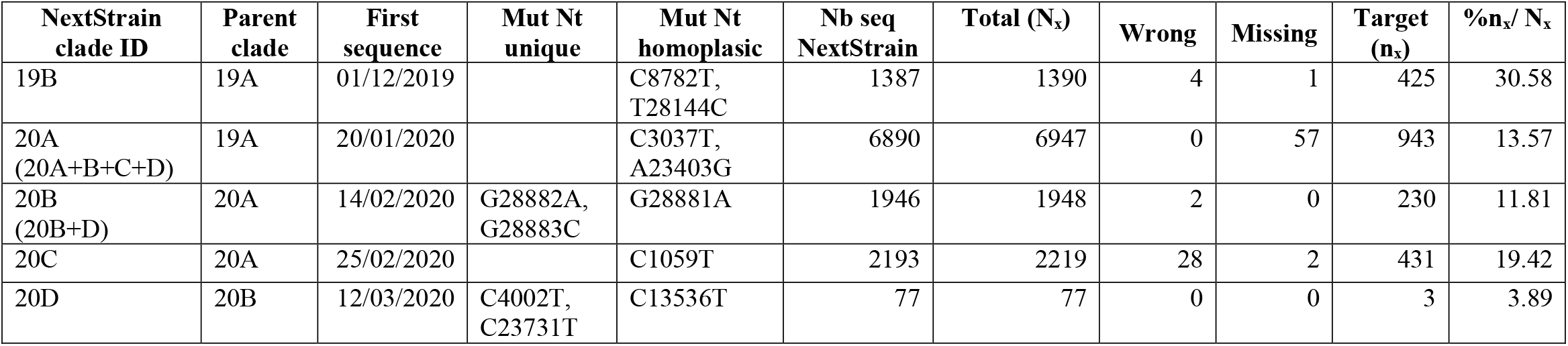
SARS-CoV-2 sampling, clade assignment and target sampling. Parent clade: parent clade of the identified clade in the first column; First sequence: date of the first sequence appearing in that clade; Mut Nt unique: nucleotide substitution and position of the unique mutations characterizing that clade (e.g., G→A at position 28,881 for clade 20B); Mut Nt homoplasic: nucleotide substitution and position of the homoplasic mutations characterizing that clade; Nb seq NextStrain: Number of sequences that were assigned to NextStrain clades; Total: size (N_X_) of the best clade in the starting reference tree (e.g., 1946+2 for the clade 20B); Wrong (*w*) and Missing (*m*) are obtained by comparing the starting reference tree with the NexStrain classification; Target: number (n_X_) of sequences of the subtype clade in the target sampling that reflects the number of cases reported by country over time during the early months of the pandemic (see Zhukova et al. 2021 for details); %n_x_ / N_X_: percentage of the number of sequences kept in the target sampling to reflect the number of cases reported by country over time during the early months of the pandemic.

| NextStrain<br>clade ID | Parent<br>clade | First<br>sequence | Mut Nt<br>unique | Mut Nt<br>homoplasic | Nb seq<br>NextStrain | Total (N<sub>x</sub>) | Wrong | Missing | Target<br>(n<sub>x</sub>) | %n<sub>x</sub>/ N<sub>x</sub> |
| --- | --- | --- | --- | --- | --- | --- | --- | --- | --- | --- |
| 19B | 19A | 01/12/2019 |  | C8782T,<br>T28144C | 1387 | 1390 | 4 | 1 | 425 | 30.58 |
| 20A<br>(20A+B+C+D) | 19A | 20/01/2020 |  | C3037T,<br>A23403G | 6890 | 6947 | 0 | 57 | 943 | 13.57 |
| 20B<br>(20B+D) | 20A | 14/02/2020 | G28882A,<br>G28883C | G28881A | 1946 | 1948 | 2 | 0 | 230 | 11.81 |
| 20C | 20A | 25/02/2020 |  | C1059T | 2193 | 2219 | 28 | 2 | 431 | 19.42 |
| 20D | 20B | 12/03/2020 | C4002T,<br>C23731T | C13536T | 77 | 77 | 0 | 0 | 3 | 3.89 |

**Figure SF2:**
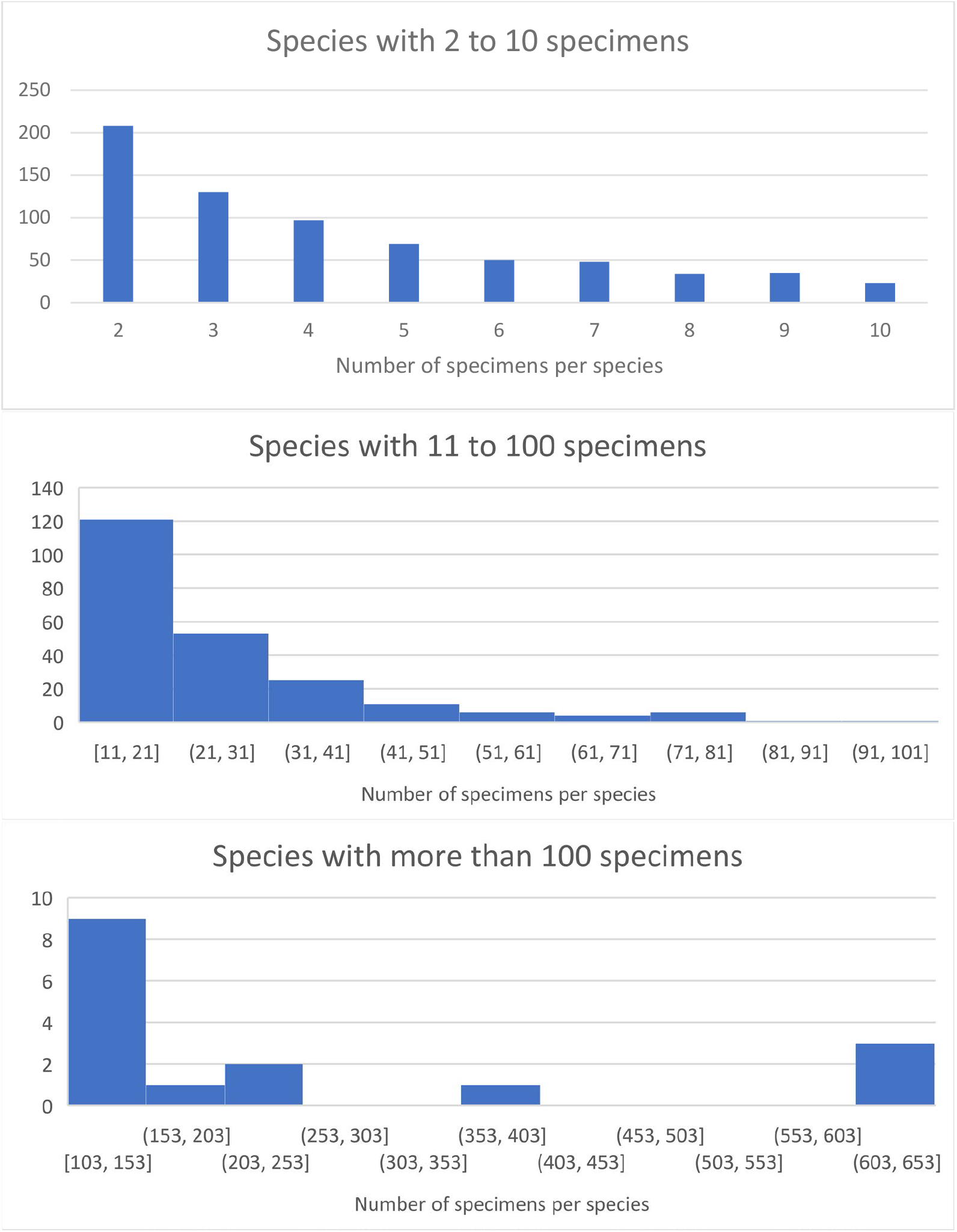
Barcode datasets: distribution of the number of specimens per delimited species. A total of 938 delimited species were used for the barcode dataset analyses. We see here that most species are represented by a few specimens (2-5) but that a few ones have extremely large sampling (> 100 barcode sequences).

